# RNA denaturation underlies circular RNA separation

**DOI:** 10.1101/2025.01.04.631262

**Authors:** Yanyi Jiang, Jørgen Kjems

## Abstract

In vitro synthesized circular RNAs (circRNAs) have emerged as a promising drug modality for RNA therapeutics due to their improved stability and reduced immunogenicity. However, effective analysis and purification of circRNAs pose critical challenges arising from the insufficient separation of circRNAs and linear RNA byproducts. In this study, we systematically evaluate the effectiveness of gel electrophoresis and high-performance liquid chromatography - size exclusion chromatography (HPLC-SEC) for separating circRNAs synthesized through ligase- or ribozyme-based strategies. While the synthesis strategy dictates the purification complexity, we demonstrate that both techniques rely on RNA denaturation to successfully separate circRNAs. Additionally, when using HPLC-SEC, we show that even a trace amount of magnesium ions in RNA samples can significantly compromise circRNA separation. Under optimized denaturing conditions, HPLC-SEC enables circRNA purification directly from diluted crude in vitro transcription products, thus streamlining the purification process. Our study provides mechanistic insights into circRNA separation, advancing the purity and scalability of circRNA-based therapeutics.

**GRAPHICAL ABSTRACT:** 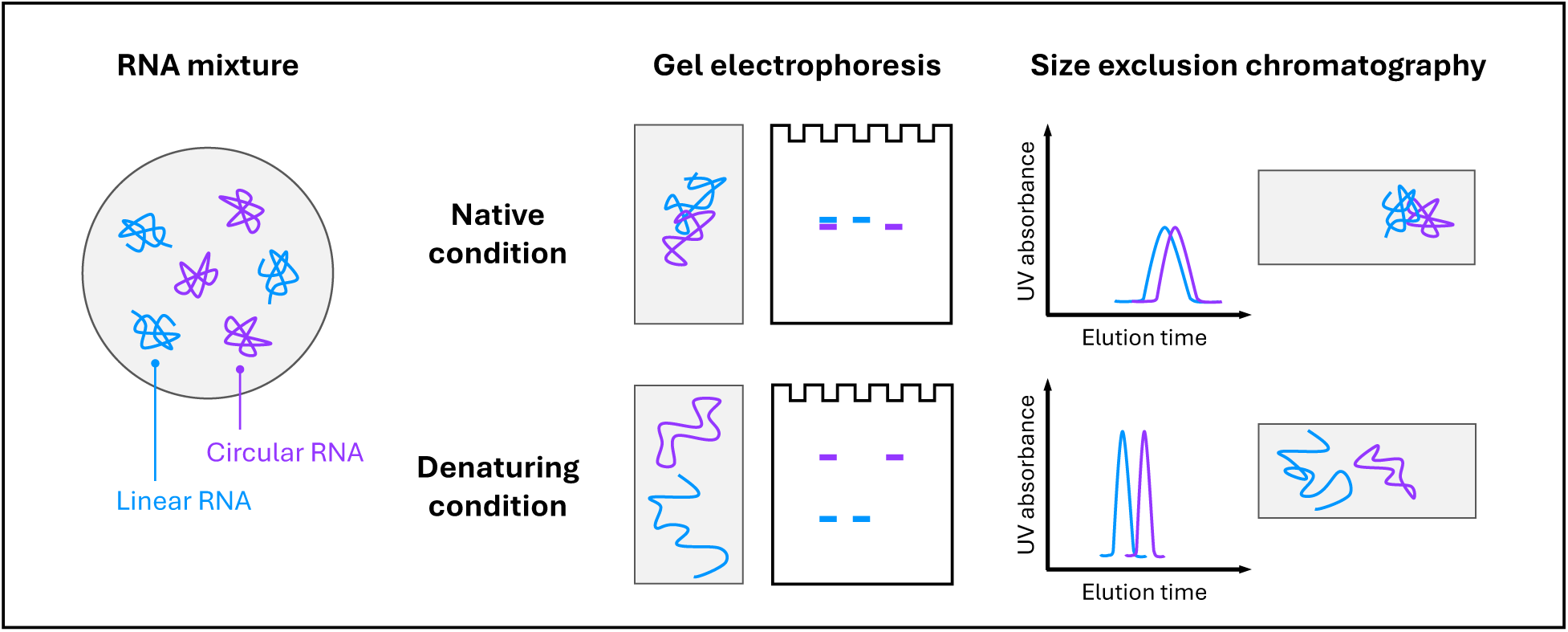

## INTRODUCTION

Following the triumph of mRNA therapeutics against COVID-19, circular RNAs (circRNAs) have garnered significant interest as a promising modality for nucleic acid therapeutics (1,2). circRNAs are single-stranded RNAs with a covalently closed loop structure, lacking free ends. This circular configuration renders circRNAs resistant to exonuclease-mediated degradation, providing them with greater stability than their linear counterparts (3). This enhanced stability makes circRNA an appealing template for robust long-term translation (4–7). Beyond protein production, the versatile therapeutic potential of circRNAs has been explored in applications such as microRNA sponges, protein sequestration, and gene editing (1). Another potent advantage of synthetic circRNAs is their reduced immunological profile. Rigorously purified, unmodified circRNA can evade cellular immune surveillance (6,8–10).

Producing in vitro synthesized circRNA entails three major steps: synthesis, purification, and analysis with quality control (Figure 1A). After the synthesis of linear RNA precursors by in vitro transcription (IVT), circularization is succeeded through intramolecular ligation using chemical, enzymatic, or ribozymatic approaches (see reviews (1,11,12); Figure 1B). Although various methods have been developed for circRNA synthesis, large-scale purification remains a significant challenge. Current purification strategies include exonuclease treatment for the selective removal of linear RNA impurities (13), phosphatase treatment to neutralize the immunogenic triphosphate group (6), and gel or HPLC purification to resolve circRNAs based on their physicochemical attributes (Figure 1A). However, the physicochemical similarity of circular and linear RNA causes them to migrate similarly when analyzed by gel electrophoresis. Given that this method serves as the cornerstone of circRNA analysis methodologies (Figure 1A), insufficient separation complicates the quality control process, potentially undermining the reliability and reproducibility of circRNA research.

**Figure 1.**
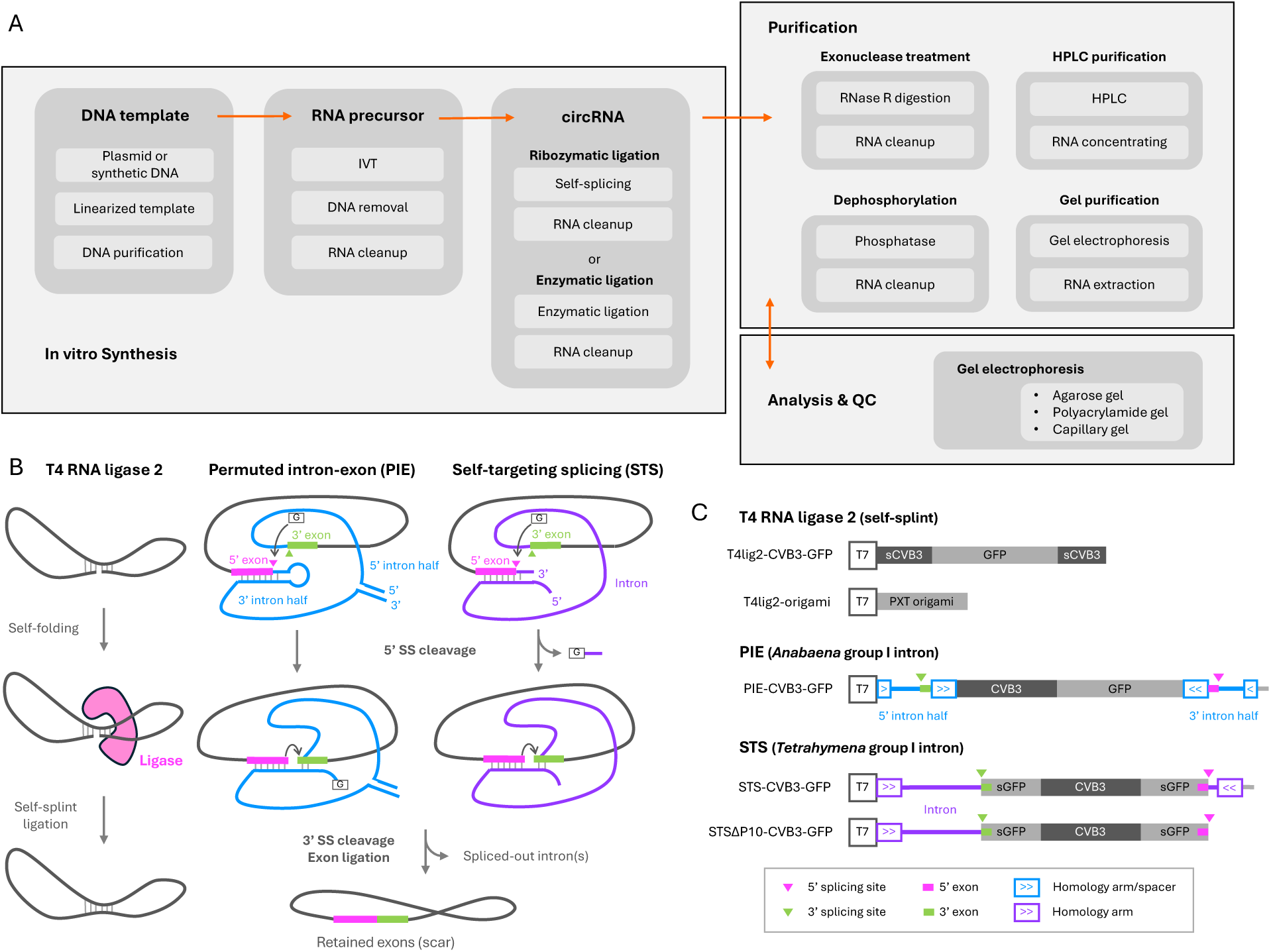
Construct design for circRNA separation investigation. (A) Workflow for producing in vitro synthesized circRNAs. (B) Mechanism of in vitro circRNA synthesis. For enzymatic circularization, T4 RNA ligase 2 recognizes a self-scaffolded double-stranded RNA precursor and seals the nick without requiring an external splint, resulting in the formation of a circRNA. For group I intron-catalyzed circularization, the 5’ and 3’ exons are ligated through self-splicing in either a cis or a trans manner, referred to as permuted intron-exon (PIE) or self-targeting splicing (STS), respectively. Both self-splicing strategies undergo two consecutive transesterification reactions, marked with arrows. In the first step, an intron-harbored exogenous guanosine cleaves the 5’ splicing site (SS). In the second step, the 5’ exon and 3’ exon are brought into proximity by conformational change and base pairing to the internal guiding sequence. The 3’-OH group of 5’ exon terminus reacts with the 3’ splicing site, resulting in ligated exons and spliced-out introns. The exon sequences are incorporated into the circRNA, forming a “scar”. G, exogenous guanosine. (C) The IVT template design in this study. The CVB3-GFP and PXT origami were the backbones for circRNA synthesis through the three methods described in (B). The CVB3 and GFP were split in the T4lig2-CVB3-GFP and STS(ΔP10)-CVB3-GFP constructs, designated as sCVB3 and sGFP, respectively. The PIE-CVB3-GFP construct utilized the original *Anabaena* exon sequences for self-splicing, in addition to homology arms and spacers to enhance the reaction, resulting in a scar of 176 nt. The STS constructs embedded exon sequences into sGFP, producing “scarless” circRNA products. The downstream sequence of the 5’ exon in the STS-CVB3-GFP construct was removed to create the STSΔP10-CVB3-GFP construct. T7, T7 promoter.

Gel electrophoresis remains the most widely used method for separating circRNAs. Depending on the gel matrix type, it includes agarose gel electrophoresis (AGE), polyacrylamide gel electrophoresis (PAGE), and capillary gel electrophoresis (CGE). In native AGE, linear and circular RNAs tend to overlap (14), while denaturing formaldehyde agarose gels, commonly used for northern blotting, facilitate circRNA separation (15). Notably, the commercial pre-cast E-Gel system allows permutated intron-exon (PIE) generated circRNAs to be separated from precursors and nicked byproducts (5,6,16), though with limited resolution compared to urea PAGE (17,18). The best circRNA separation resolution is achieved using denaturing urea PAGE, where circRNAs migrate distinctly slower than their linear counterparts (19). In contrast, native PAGE provides poor separation efficiency for circRNAs (20,21). CGE, commonly applied for rapid RNA quality assessment, is also employed for circRNA quality evaluation (18,22–27). While this technique can distinguish PIE-generated circRNAs from their longer precursors, its ability to separate circRNAs from linear RNA counterparts of the same size remains controversial (28). Among these three electrophoresis methods, both AGE and PAGE can serve for circRNA purification (5,8), but the inefficiency of the downstream gel extraction of the circRNA limits their application for large-quantity manufacturing.

In contrast, high-performance liquid chromatography - size exclusion chromatography (HPLC-SEC) offers scalability for the purification of biopharmaceutical substances (29) and has been utilized for circRNA purification since 2018 (5). In HPLC-SEC, circRNAs elute later than their linear counterparts, allowing for their separation (5). However, the separation remains incomplete as the circRNA fraction partially overlaps with the linear fraction. To obtain relatively pure circRNA, the latter portion of the circRNA fraction is typically collected, although this approach compromises the yield (6,7,30,31). Factors influencing SEC performance in circRNA purification have not been systematically studied but may include column particle pore size, circRNA size, circRNA synthesis method, and mobile phase conditions (reviewed in Table 1). Of note, the choice of analysis and quality control methodology directly impacts the purity assessment of circRNAs.

**Table 1.** Literature summary of circRNA HPLC-SEC purification. PIE: permuted intron-exon. TRIC: *trans*-ribozyme-based circularization. Ana: *Anabaena* pre-tRNA group I intron. td: bacteriophage T4 thymidylate synthase gene group I intron. Cte: *Clostridium tetani* group II intron. TE: Tris-EDTA buffer, 10 mM Tris-HCl, 1 mM EDTA. PB: Phosphate buffer with variable ratio of NaH_2_PO_4_ and Na_2_HPO_4_. AGE: agarose gel electrophoresis. PAGE: polyacrylamide gel electrophoresis. CGE: capillary gel electrophoresis. RP-HPLC: reverse-phase HPLC.

| Publication Year | circRNA Synthesis | Approximate circRNA Size (nt) | SEC Column (brand, pore size, particle size) | Mobile Phase | circRNA Analysis | Literature Source |
| --- | --- | --- | --- | --- | --- | --- |
| 2018 | PIE-Ana | 1500 | Sepax, 2000 Å, 5 µm | TE, pH 6.0 | AGE (E-Gel) | (5) |
| 2019 | PIE-Ana | 1500 | Sepax, 2000 Å, 5 µm | TE, pH 6.0 | AGE (E-Gel) | (6) |
| 2019 | PIE-td | 1500 | Sepax, 2000 Å, 5 µm | TE | AGE, CGE | (32) |
| 2020 | PIE-Ana | 350 to 500 | Sepax, 2000 Å, 5 µm | TE, pH 6.8 | AGE (E-Gel) | (30) |
| 2022 | PIE-Ana<br>T4 RNA ligase 2 | 1600 to 1900 | Sepax, 2000 Å, 5 µm | TE, pH 7.5 | AGE, urea PAGE | (7) |
| 2022 | T4 RNA ligase 1<br>T4 RNA ligase 2 | 150 | Sepax, 2000 Å, 5 µm | TE | AGE | (33) |
| 2022 | PIE-Ana | 1100 to 1400 | Sepax, 2000 Å, 5 µm | TE, pH 6.0, at 35°C | AGE (E-Gel) | (16) |
| 2022 | PIE-Ana, PIE-td<br>T4 RNA ligase 1 | 300 to 550 | Sepax, 2000 Å, 5 µm | TE, pH 6.0 | AGE, urea PAGE | (34) |
| 2022 | PIE-Cte<br>(group II intron) | 1500 | Waters XBridge,<br>450 Å, 3.5 µm | TE + PB, pH 7.4, at 25°C | AGE | (35) |
|  |  |  | Sepax, 1000 Å, 5 µm | TE + PB, pH 7.4, at 25°C | AGE |  |
| 2022 | PIE-Ana | 1800 | Sepax, 1000 Å, 5 µm | PB, pH 6.0 | AGE, CGE | (36) |
| 2022 | PIE-Ana | 1700 to 2700 | Sepax, 2000 Å, 5 µm | TE, pH 6.0 | AGE, CGE | (31) |
| 2024 | PIE-Ana | 1500 | Sepax, 2000 Å, 5 µm | Not specified | AGE | (37) |
| 2024 | PIE-Ana | 2700 | Sepax, 2000 Å, 5 µm | TE, pH 6.0 | AGE | (38) |
| 2024 | PIE-Ana | 1600 | Sepax, 1000 Å, 5 µm | PB, pH 7.0 | CGE, RP-HPLC | (28) |
| 2024 | PIE-Ana, PIE-td | 1500 | Sepax, 1000 Å, 5 µm | PB, pH 7.0 | AGE | (39) |
| 2024 | PIE-Ana | 1600 | Sepax, 1000 Å, 5 µm | PB | CGE | (40) |
| 2024 | TRIC-Ana | 1400 | Sepax, 2000 Å, 5 µm | TE, pH 6.5 | (Urea) AGE | (10) |

In this study, we have conducted a comprehensive investigation to evaluate the separation effectiveness of circRNA using gel electrophoresis and HPLC-SEC, revealing that RNA denaturation is critical for effective circRNA separation. Comparing circRNAs synthesized through enzymatic and ribozymatic methods, we highlight the purification complexities arising from the differing byproduct profiles and the sample processing associated with each synthesis method.

## MATERIALS AND METHODS

### In vitro transcription (IVT)

The sequences of PIE-, STS-, and self-splint T4lig2-CVB3-GFP were obtained from previous published studies (5,7,41). The DNA templates were synthesized by Integrated DNA Technologies or Twist Bioscience (sequences shown in Table S1). The IVT templates were linearized and amplified through PCR using Q5 DNA polymerase (NEB), followed by agarose gel purification using Monarch DNA Gel Extraction Kit (NEB). RNAs were in vitro transcribed using the HiScribe T7 Quick High Yield RNA Synthesis Kit (NEB). Specifically, 100-500 ng purified DNA template was added into 20 μL reaction volume and incubated at 37°C overnight. To produce the RNA precursor with 5’ monophosphate for T4 ligation, additional GMP (Sigma) was added (50 mM final concentration) to the IVT reaction. After incubation, the IVT sample was diluted to 50 µL using RNase-free water and treated with 2 µL DNase I (NEB) at 37°C for 15 minutes. The RNA was subsequently column-purified using the MEGAclear Transcription Clean-Up Kit (Invitrogen) according to the manufacturer’s instructions.

### RNA circularization

For T4 ligation constructs, transcribed linear RNA precursors were incubated with T4 RNA ligase 2 (NEB) following the manufacturer’s protocol at 37°C for 2 hours. For the PIE construct, 5 µL 10x T4 RNA ligase buffer (500 mM Tris-HCl, 100 mM MgCl_2_, 10 mM DTT, pH 7.5; NEB) and GTP (final concentration of 2 mM) were added to column-purified RNA samples, bringing the total volume to 50 µL, followed by incubation at 55°C for 15 to 30 minutes. After the circularization process, RNAs were column-purified by RNA Clean & Concentrator-25 (Zymo Research) for subsequent studies. For STS constructs, no circularization step was performed after DNase I treatment (17).

### Enzymatic purification of circRNA

RNase R (Biosearch Technologies) treatment was performed after RNA cleanup at 1 U/µg RNA, with incubation at 37°C for 15 to 60 minutes. In the control groups, the RNase R was replaced by RNase-free water. Since the performance of RNase R can vary between batches and RNA constructs, conducting a pilot test and optimizing incubation time and enzyme concentration are necessary. Following RNase R treatment, RNA was directly subjected to electrophoresis analysis without column purification.

### Agarose and polyacrylamide gel electrophoresis

Self-cast agarose gels were prepared by dissolving 1% or 2% (w/v) agarose (UltraPure, Invitrogen) in 1x TBE buffer (100 mM Tris, 90 mM boric acid, and 1 mM EDTA, pH 8.3), diluted from UltraPure TBE 10x stock (Invitrogen) with ultrapure water. Before gel solidification, 3-5 µL SYBR Safe (Invitrogen) was pre-mixed into 100 µL of the melted gel. RNA samples were mixed with a denaturing 2x RNA Loading Dye (95% formamide, 0.02% SDS, 0.02% bromophenol blue, 0.01% xylene cyanol, 1 mM EDTA; NEB), followed by snap cooling (90°C for 1 minute, then on ice for at least 5 minutes). The gel was run in 1x TBE buffer using 150 to 200 V at room temperature.

For the E-Gel electrophoresis system, three types of E-Gels (Invitrogen) were used: 1% non-EX gel (A45203), 2% non-EX gel (A45205), and 2% EX gel (G402022). RNA samples (50-1000 ng) were diluted to 20 μL with nuclease-free water and loaded onto pre-cast gels without prior snap-cooling. Electrophoresis was conducted using E-Gel Power Snap Electrophoresis Device (Invitrogen), following either the “E-Gel 1-2% EX” program for 20 minutes or the “E-Gel 0.8-2%” program for 20 to 40 minutes. The temperatures of E-Gels were measured immediately after electrophoresis using a miniature infrared thermometer (Elma 608, Elma Instruments).

For denaturing PAGE, urea polyacrylamide gels were prepared using the UreaGel System Kit (National Diagnostics), where UreaGel concentrate (19:1 acrylamide/bisacrylamide, 7.5 M urea) was mixed with UreaGel diluent (7.5 M urea) at varying ratios depending on the desired gel percentage, and TBE buffer (1x final concentration, pH 8.3). For polymerization, 80 µL 10% (w/v) APS and 4 µL TEMED were added to a 10 mL gel solution. RNA samples or HPLC fractions were mixed with the denaturing 2x RNA Loading Dye (NEB), followed by snap cooling (90°C for 2 minutes, then placed on ice for at least 5 minutes). Electrophoresis was performed in 1x TBE buffer at 350 V and 20 W for 20-30 minutes at room temperature.

For native PAGE, AccuGel premixed solution (40% (w/v), 29:1 acrylamide/bisacrylamide, National Diagnostics) was diluted with ultrapure water and TBE buffer (1x final concentration, pH 8.3) to prepare the gel. For polymerization, 80 µL 10% (w/v) APS and 4 µL TEMED were added to a 10 mL gel solution. RNA samples were combined with a non-denaturing 6x loading dye (TrackIt Cyan/Yellow Loading Buffer, 10 mM Tris-HCl, 60 mM EDTA, 0.03% xylene cyanol FF, 0.3% tartrazine, 60% glycerol, pH 7.6; Invitrogen) before gel loading. Electrophoresis was carried out in a 4°C cold room using 1x TBE as the running buffer at 80 V for 130 minutes.

All polyacrylamide gels were prepared using mini gel cassettes (8 x 8 cm, 1.0 mm, Invitrogen), and the electrophoresis was performed using XCell SureLock Mini-Cell (Invitrogen). After electrophoresis, the polyacrylamide gels were stained with SYBR Gold (Invitrogen) to visualize RNA bands. The ssRNA ladder (NEB) was used as the size marker. Both agarose and polyacrylamide gels were visualized by an Amersham Typhoon biomolecular imager (Cytiva).

### Capillary gel electrophoresis

RNA samples (1 µL) were loaded onto a microfluidic chip from the RNA 6000 Nano Kit (Agilent, 5067-1511) following the manufacturer’s instruction. For the pre-denaturing condition, before loading, RNA samples were mixed with the denaturing 2x RNA Loading Dye (NEB), heated at 65°C for 2 minutes, and snap-cooled on ice. The samples were analyzed using the 2100 Bioanalyzer (Agilent).

### HPLC-SEC

RNA samples (1-5 µg) were injected to a size exclusion column on an Agilent 1100 Series HPLC (Agilent), with RNase-free TE buffer (10 mM Tris, 1 mM EDTA, pH 7.5; 20x stock, Thermo Fisher) as mobile phase at a flow rate of 0.3 mL/minute (SEC-1000/2000) or 0.7 mL/minute (SEC-4000). The SEC columns are with particle size of 5 μm and pore size of 2000 Å (4.6 x 300 mm; Sepax, 215980P-4630) or 1000 Å (4.6 x 300 mm; Sepax, 215950P-4630) or 4000 Å (8 x 300 mm; Altmann Analytik, AANRS4SE-5030080). The SEC-2000 and SEC-1000 columns were wrapped with copper wires and connected to an extended stainless steel capillary (0.17 x 450 mm, Agilent) in the inlet. The capillary was attached to a heating plate to preheat the mobile phase before it entered the column (Figure S1A) (42). RNA was detected by UV absorbance at 260 nm. The collected HPLC fractions were analyzed by 2% E-Gels or 3.5% urea PAGE. The separation resolution (R_s_) was calculated using the following equation,

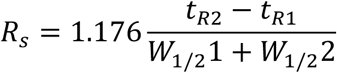

where *t_R1_* and *t_R2_* are the retention times of the two peaks, and *W*_1/2_1 and *W*_1/2_2 represent the peak widths at half height for each peak.

## RESULTS

### Analysis of circRNAs with gel electrophoresis

To investigate how circRNA synthesis methods affect the subsequent analysis in gel electrophoresis, we designed five constructs where two different circRNAs are produced using three different circularization methods (Figure 1C). Three of the constructs contain the commonly utilized coxsackievirus B3 (CVB3) internal ribosome entry site (IRES) upstream of GFP open reading frame: 1) T4 RNA ligase 2 mediated self-splint ligation (T4lig2-CVB3-GFP) (7,43); 2) permuted intron-exon (PIE) *Anabaena* group I intron mediated circularized through cis self-splicing (PIE-CVB3-GFP) (5,44); and 3) self-targeting splicing (STS) *Tetrahymena* group I intron mediated circularization (STS-CVB3-GFP) (17,41). A variant of the latter construct where the downstream sequence of the 5’ splicing site was deleted was also included (STSβ10-CVB3-GFP). It exhibits higher RNA circularization efficiency, likely due to the omission of the first esterification reaction leading to 5’ splicing site cleavage (Figure 1B) (17). As an example of a shorter (415 nt) circRNA without translation elements, we used the paranemic crossover triangle (PXT) RNA origami (45) adopted for T4 RNA ligase 2 mediated self-splint ligation (T4lig2-origami).

The product profile differs depending on circRNA synthesis strategies. T4 enzymatic ligation primarily produces unreacted precursors of the same size as circRNAs, while ribozyme-catalyzed circularization involves longer precursors, potential splicing intermediates, and excised introns (Figure 1B). Common to all strategies, inevitable circRNA nicking leads to linear side-products, either matching the size of circRNAs or shorter if multiple nicks occur.

Native agarose gel is commonly employed to analyze synthetic circRNAs. Here, we investigated the ability of 1% and 2% self-cast and commercial pre-cast agarose E-Gels to separate circRNA from linear contaminates (Figure 2). For all reactions, the exonuclease RNase R was used to identify the bands corresponding to circRNAs, as it selectively digests the linear RNA with a free 3’ end while leaving circRNA intact (13). Self-cast 1% agarose gels were unable to separate circular and linear CVB-GFP RNAs of the same size (Figure 2B). In 2% agarose gel, circRNA migrates slightly faster than its linear counterpart, but the resolution is very limited, as the linear and circRNA bands remain close, even after extended electrophoresis (Figure 2C). The commercial pre-cast agarose E-Gel has been reported to separate circRNA from its linear counterpart (5,6,16). There are two types of E-Gels, EX and non-EX, which can be run using various electrophoresis programs. Using an EX 1-2% electrophoresis program for 20 minutes, both 2% EX and 2% non-EX gels could separate circRNAs from linear ones, but 1% non-EX gel could not (Figure 2D). Using a non-EX 0.8-2% electrophoresis program, circRNA could be separated by two types of 2% E-Gels but with lower resolution, and non-EX gel required extended electrophoresis to achieve circRNA separation (Figure 2E). Notably, after electrophoresis, the gel temperature differed among programs and E-Gel types. EX and non-EX gels reached above 80°C with EX 1-2% program after 20-minute electrophoresis, whereas the gel temperature only reached approximately 55°C with non-EX 0.8-2% program for 24 to 40 minutes, indicating higher temperature may facilitate circRNA separation.

**Figure 2.**
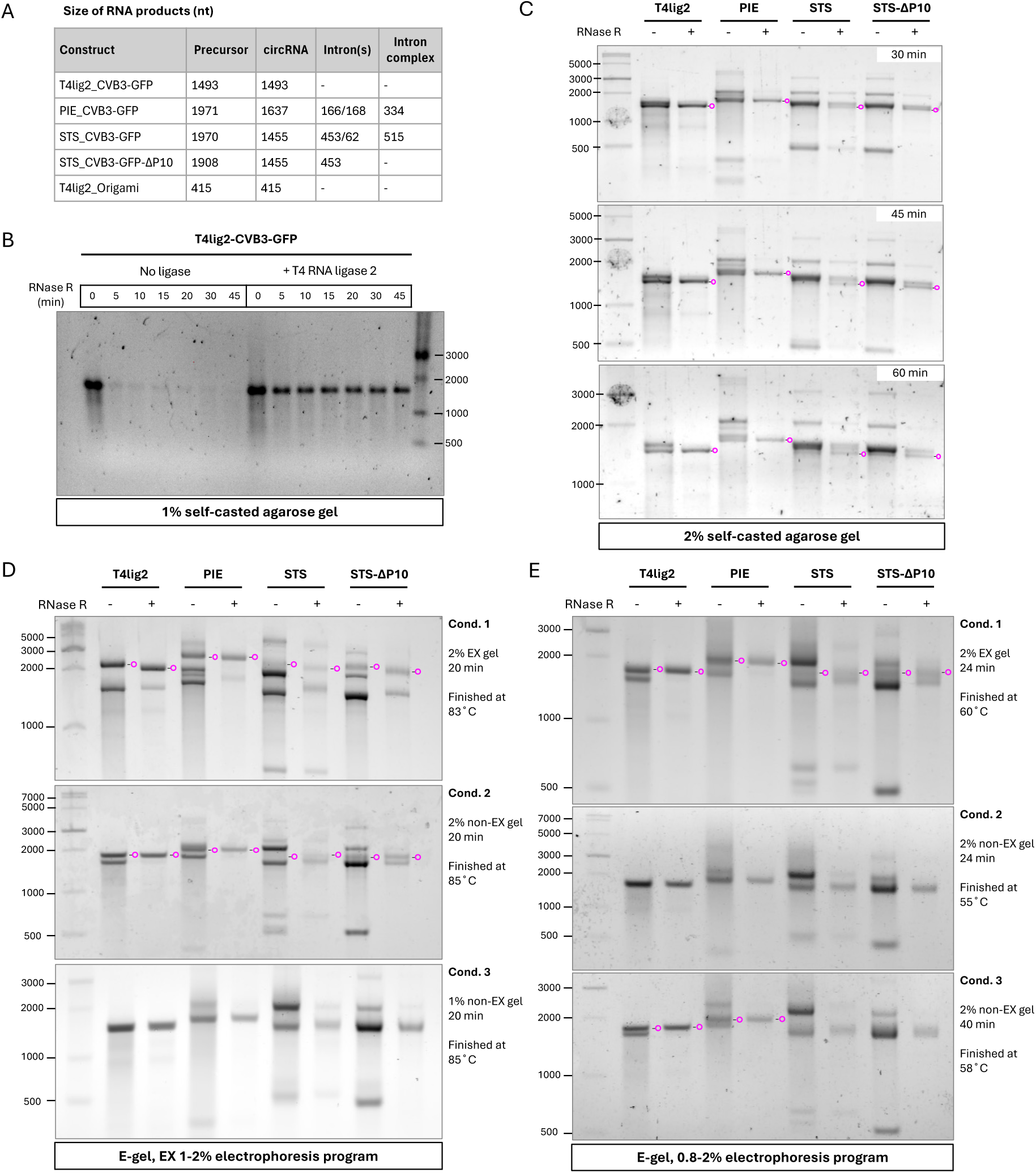
Agarose gel analysis of circRNAs. **(A)** The predicted size of RNA products formed upon the circularization of the different constructs. **(B)** Analysis of products in a 1% self-casted agarose gel. The RNA precursor from T4lig2-CVB3-GFP was incubated with or without T4 RNA ligase 2, followed by RNase R treatment (0 to 45 minutes) and subsequent gel electrophoresis. **(C)** Analysis of products in a 2% self-casted agarose gel. The four denoted RNA constructs were treated with RNase R and analyzed using the indicated electrophoresis running times. **(D)** E-Gel analysis of circRNAs in 1% or 2% E-Gels using the “E-Gel EX 1-2%” program. The four denoted RNA constructs were treated with or without RNase R before loading. The surface temperatures of E-Gels were measured immediately after the electrophoresis was done. **(E)** E-Gel analysis of the same samples as in (D) but using the “E-Gel 0.8-2%” program. The bands corresponding to circRNAs are denoted with magenta circles.

Denaturing urea polyacrylamide gel electrophoresis (PAGE) provides superior circRNA separation efficacy where circRNAs migrate markedly slower than linear ones (19). In contrast, native PAGE has difficulties separating linear and circular RNAs, with circRNAs either migrating similarly or slightly faster than their linear forms (20,21). First, we investigated how gel conditions and sample pre-treatment affect the separation of CVB3-GFP and PXT origami derived circRNAs (Figure 3A). Given that the linear RNA precursor of T4lig2-origami resisted RNase R digestion because of its compact 3’ end structure (46), we chose the T4 ligase-untreated group as the linear control. For native 3.5% PAGE performed at 4°C, circular T4lig2-origami (415 nt) migrated slightly faster than its linear form, which aligns with the former finding (21) (Figure 3A, Condition 1). Under native condition, the circRNA separation of CVB3-GFP constructs (1450-1650 nt) was poor, and smearing circRNA bands were observed in three ribozyme-derived groups (Figure 3A, Condition 1). Multiple RNA conformation may co-exist in the CVB3-GFP circRNAs, while T4lig2-origami appeared as a sharper band in the native gel analysis, probably due to its smaller size and lower structural complexity. When the PAGE was processed at room temperature, circRNAs moved slower than linear counterparts in the gel (Figure 3A, Condition 2). Pre-denatured samples with 47.5% formamide and snap cooling before gel loading further increased separation resolution (Figure 3A, Condition 3). In 7.5 M urea denaturing PAGE, circRNAs migrated even slower, and smearing bands appearing in native gels were sharpened, resulting in improved separation efficiency (Figure 3A, Condition 4). These results demonstrate how RNA denaturation, in general, improves circRNA separation from linear counterparts. Next, we tested circRNA migration under varying gel concentrations in denaturing PAGE (Figure 3B). Reducing the gel concentration from 4% to 2.5% lowered the separation efficiency for circRNAs. In 2.5% denaturing PAGE, STS(ΔP10)-CVB3-GFP circRNAs even migrated faster than the linear precursors. Considering Reducing the gel concentration from 4% to 2.5% reduced the separation efficiency for circRNAs. In 2.5% denaturing PAGE, circRNAs migrated faster than their linear precursors in the STS constructs. Considering both electrophoresis time and separation efficiency, we selected 3.5% urea PAGE for the subsequent analysis of PIE and STS constructs.

**Figure 3.**
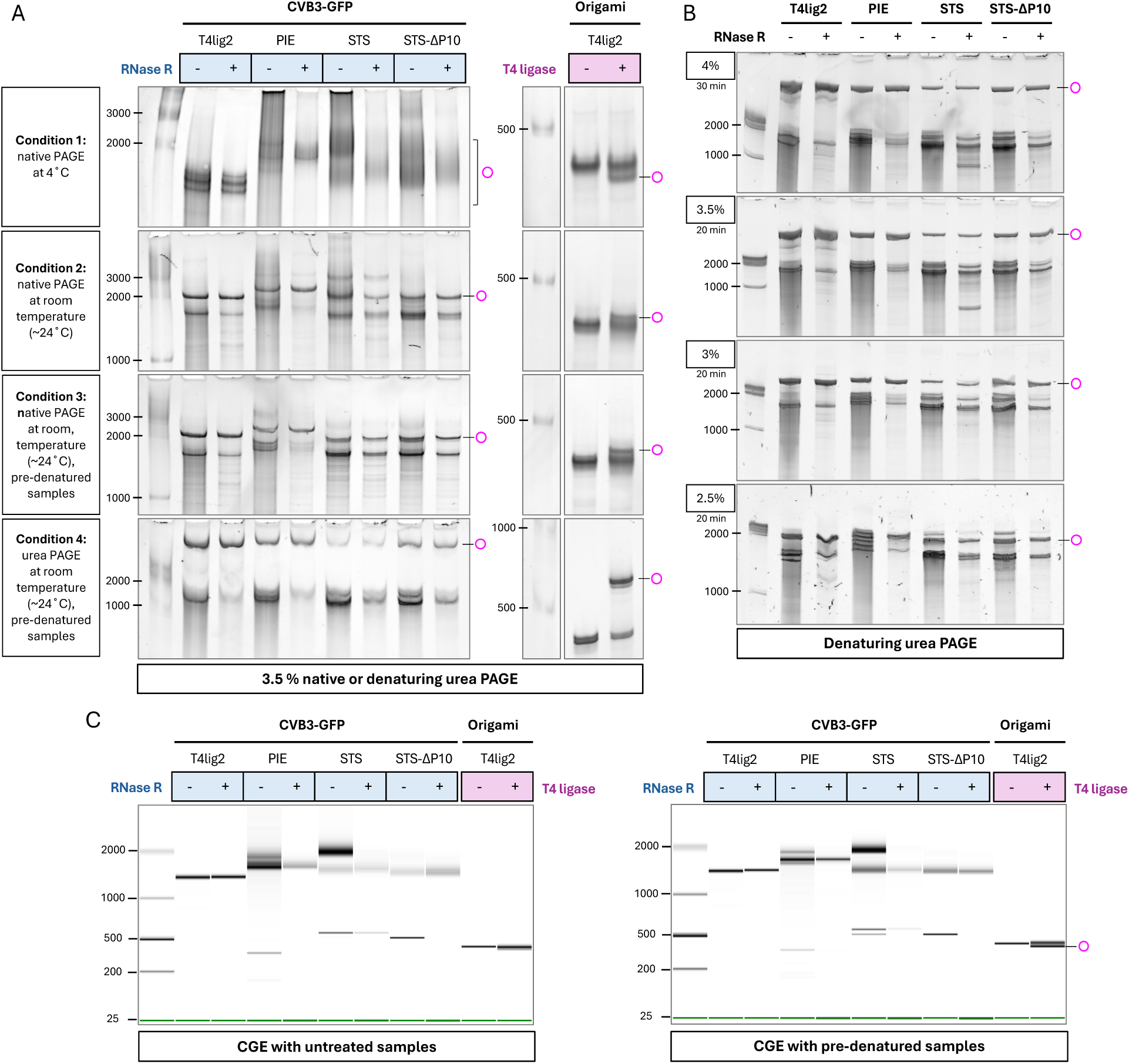
PAGE and CGE analysis of circRNAs. **(A)** PAGE analysis of circRNAs under native and denaturing conditions. The CVB3-GFP circRNA samples (left panel) were incubated with or without RNase R. The origami RNA precursor (right panel) was incubated with or without T4 RNA ligase 2. Four different PAGE conditions were tested: (1) Native PAGE. RNA samples were mixed with native loading dye and analyzed in a 3.5% native polyacrylamide gel at 4°C (80 V for 130 minutes). (2) Same as Condition 1, but run at room temperature (350 V for 16 minutes). (3) Same as Condition 2 but with RNA samples pre-denatured by mixing with denaturing 2x RNA Loading Dye (NEB), heating at 90°C for 2 minutes, and then snap-cooling on ice before being analyzed. (4) Same as Condition 3 but using denaturing PAGE (7.5 M urea). **(B)** Denaturing PAGE analysis of circRNAs in gels at indicated acrylamide concentrations. Pre-denatured RNA samples were analyzed by 7.5 M urea PAGE at 350 V for 20 or 30 minutes at room temperature. **(C)** Capillary gel electrophoresis of circRNAs. Products from the denoted RNA constructs were analyzed using the 2100 Bioanalyzer (Agilent). For the pre-denaturing condition (right), RNAs were mixed with 2x denaturing RNA loading dye, heated at 65°C for 2 minutes, and then snap-cooled on ice prior to chip loading. The bands corresponding to circRNAs are marked with magenta circles to the right.

Next, we investigated the circRNA separation using capillary gel electrophoresis (CGE) with and without pre-denaturing the samples before loading (Figure 3C). Under native condition, CGE failed to separate the T4lig2-CVB3-GFP and T4lig2-origami circRNAs when the circRNAs share the same size as their linear precursors (Figure 3C, left panel). However, when samples were pre-denatured with formamide and snap cooling, the circRNA from T4lig2-origami exhibited slightly faster migration than the linear precursor (Figure 3C, right panel). Additionally, we observed that pre-denaturation resulted in sharper bands in CGE, possibly attributed to increased homogeneity for RNA configuration following the denaturing process. Since the circRNA yield was quite low from the STS-CVB3-GFP construct (Figure 3B), we opted for STSΔP10-CVB3-GFP in our subsequent studies.

### Column heating enables the effective separation of circRNAs in HPLC-SEC

Inspired by the observation that RNA denaturation during electrophoresis is crucial for effective circRNA separation, we hypothesized that this principle can be utilized in HPLC-SEC. When heating the SEC column, we noticed that the air temperature of the HPLC column oven was lower than the setting (Figure S1A). To ensure the SEC column constantly stayed at the pre-set temperature, we wrapped the column with a copper wire to facilitate better heat conduction and attached an extended tubing to the heating plate to preheat the mobile phase before it entered the column (42) (Figure S1A).

We first tested the circRNA separation resolution (R_s_) in SEC-2000 (2000 Å pore size, 5 µm particle size) using the T4lig2-CVB3-GFP construct and analyzed the elution fractions by E-Gel electrophoresis (Figure 4A). The resolution was measured by both retention time and peak width at half height (see Methods). We found that the circRNA peak largely overlapped with the precursor peak at a column temperature of 30°C. However, heating the column to 45°C or 60°C significantly improved the resolution. Column heating also decreased the retention times for both linear and circular RNA fractions, with a more pronounced effect on linear RNAs, resulting in overall better resolution. Next, we tested SEC-1000 (1000 Å pore size, 5 µm particle size) using the same samples and analyzed the eluted fractions by E-Gel electrophoresis. Heating of the SEC-1000 column also resulted in a better circRNA separation, but the resolution was poorer than using the SEC-2000 column (Figure 4A).

**Figure 4.**
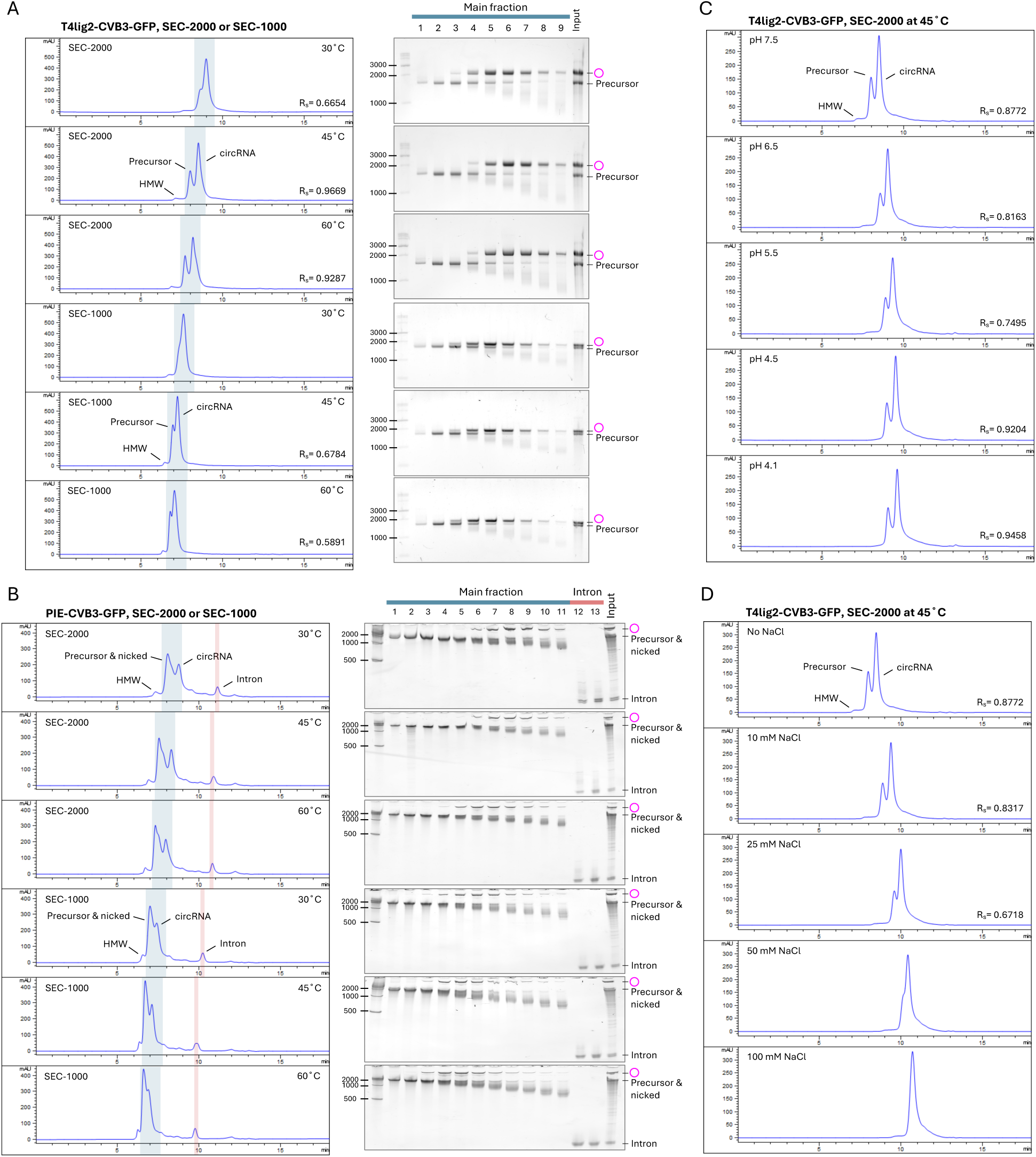
HPLC-SEC purification of circRNAs. **(A)** Separation of T4lig2-CVB3-GFP-derived products in SEC-1000/2000 at different column temperatures. RNAs were purified with SEC-1000 or -2000 at 30, 45 and 60°C. The eluted fractions (marked in blue) were analyzed using E-Gels (2% EX gel for SEC-2000 elutes, 2% non-EX gel for SEC-1000 elutes). HMW, high molecular weight. **(B)** Same as in (A) but analyzing the separation of PIE-CVB3-GFP-derived products, by 3.5% urea PAGE at the indicated temperature. Intron fractions are depicted in red. **(C)** T4lig2-CVB3-GFP derived RNA separated by the SEC-2000 column at 45°C, with mobile phases adjusted to the indicated pH. **(D)** T4lig2-CVB3-GFP-derived RNA was purified using the SEC-2000 column at 45°C with mobile phases containing the indicated salt concentrations.

As expected, the HPLC chromatographic profiles of PIE and STS constructs appeared more complicated than those of the T4 ligation system. The separation of PIE-CVB3-GFP circRNAs was also enhanced when the column was heated to 45°C according to the analysis of the fractions by urea PAGE (Figure 4B). For the STSΔP10-CVB3-GFP construct, we observed a plethora of nicked or non-circularized RNAs. Heating the column from 30°C to 45°C did not significantly increase the resolution, and there was a large overlap of the circRNA and linear peaks at 60°C (Figure S1B). Interestingly, for the STSΔP10-CVB3-GFP construct, spliced-out introns co-eluted with precursor and circRNA fractions at 30°C and 45°C, but were separated at 60°C. Again, SEC-2000 outperformed SEC-1000 in separating the circRNA in both PIE and STS systems, consistent with previous observations from the T4 ligation system. We hypothesized larger pore size might facilitate circRNA separation and tested SEC-4000 (4000 Å pore size, 5 µm particle size). However, SEC-4000 failed to differentiate circRNA from precursors and small introns at both 30°C and 65°C (Figure S2).

### pH and salt in the SEC mobile phase influence circRNA separation

A comprehensive study of circRNA separation in different SEC mobile phases is currently lacking. Therefore, we evaluated the resolution of circRNA separation using a Tris-EDTA buffer with a pH range from 7.5 to 4.1, utilizing SEC-2000 column heated to 45°C. For the T4lig2-CVB3-GFP construct, we observed a decline in resolution when lowering the pH from 7.5 to 5.5, but a significant improvement when lowering the pH further from 5.5 to 4.1 (Figure 4C). A similar trend was seen for the PIE-CVB3-GFP construct (Figure S3A), whereas the difference was not discernible for the STSΔP10-CVB3-GFP construct (Figure S3B).

We then tested how salt concentrations in the SEC mobile phase affect circRNA separation. For the T4lig-CVB3-GFP RNA injected into a SEC-2000 column at 45°C, increasing the sodium chloride concentrations from 0 to 100 mM had a negative effect on circRNA separation efficiency (Figure 4D). In contrast, for the PIE and STS constructs, low salt conditions (10 mM NaCl) in the mobile phase slightly improved circRNA separation (Figure S4). However, at higher salt concentrations (> 25 mM NaCl), the SEC could not separate the circRNAs for any of the constructs.

### Pre-treatment of RNA affects circRNA separation in HPLC-SEC

An RNA cleanup is typically required prior to HPLC injection to remove small molecules, such as salts and unincorporated nucleotides, as well as proteins derived from enzymatic reactions (Figure 1C). We hypothesized that subtle differences in the stringency of the RNA cleanup prior to HPLC injection could affect the chromatographic profiles. Therefore, we investigated how sample treatment and residual materials affect SEC performance for circRNA separation (Figure 5, S5). We first tested snap cooling of RNA at 65°C before injecting it into the HPLC system. This resulted in a slight increase in resolution for the T4lig2-CVB3-GFP construct, as both circRNA and precursor peaks became sharper (Figure 5A, Condition 2). Interestingly, the snap cooling process diminished the co-elution of introns with larger RNA species for both PIE and STS constructs (Figure 5B, S5, Condition 2). Sufficient magnesium ion concentration is critical for IVT and ribozyme activity. Thus, we examined how adding magnesium to spin column-purified RNA samples affects SEC performance. The separation of circRNAs, derived from T4 ligation, PIE, and STS constructs, was compromised by the presence of 10 mM magnesium in RNA samples (Figure 5A-B, S5, Condition 3). Additionally, more introns co-fractionated with precursors and circRNAs, and more circRNAs were detected in precursor fractions. When snap cooling was applied to magnesium-containing RNA samples, the subsequent HPLC-SEC could not separate circRNAs (Figure 5A-B, S5, Condition 4). In contrast, the addition of 10 mM EDTA, a strong chelating agent that deactivates divalent metal ions, followed by snap cooling mitigated the issue of intron co-elution in STSΔP10-CVB3-GFP circRNA fractions, resulting in improved circRNA purification (Figure S5, Condition 5).

**Figure 5.**
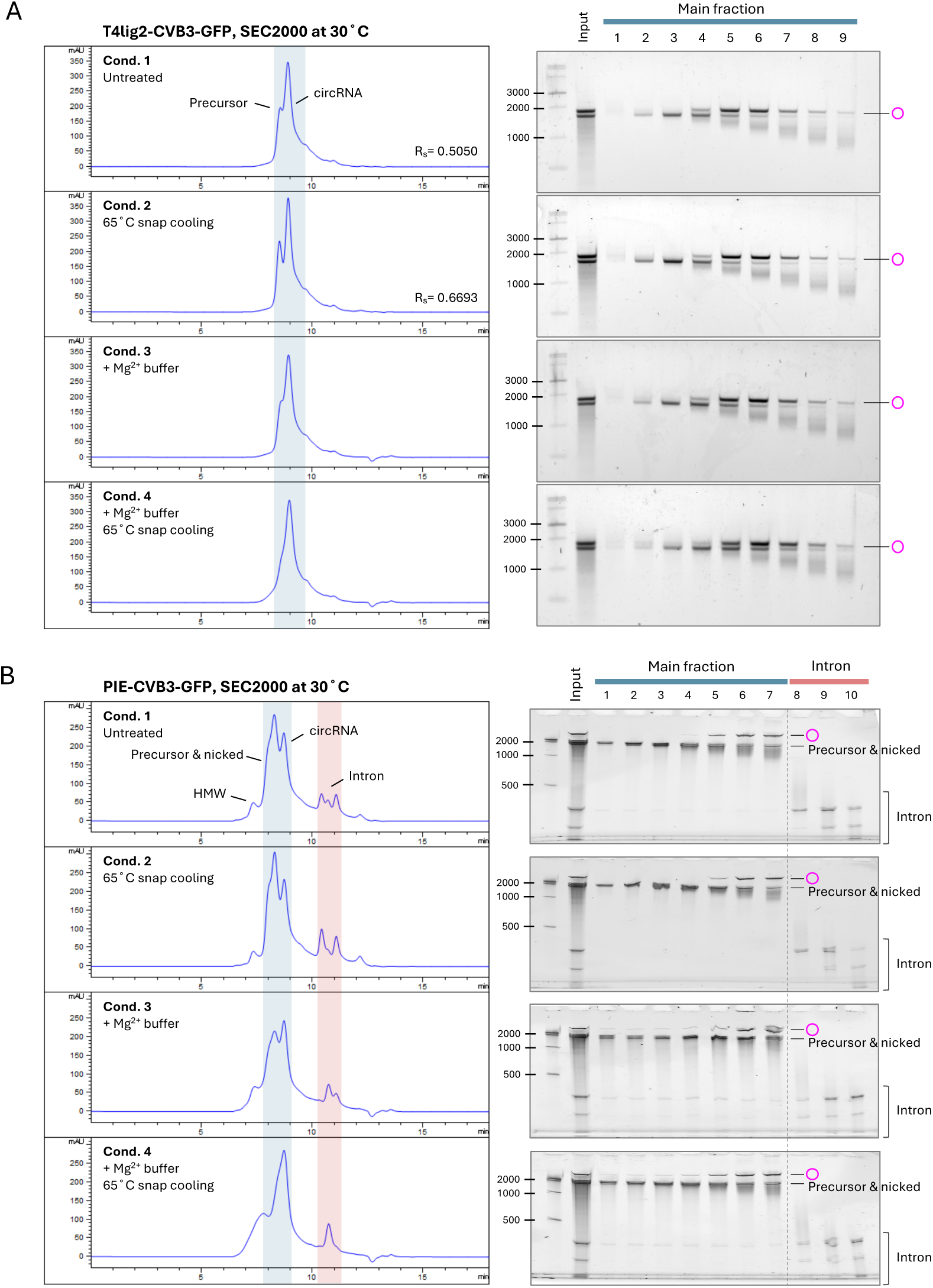
Effect of RNA pre-treatment on circRNA separation in HPLC-SEC. **(A)** Effect of the indicated pre-treatment of the T4lig1-CVB3-GFP RNAs prior to HPLC-SEC purification. Processed samples were subsequently injected into the SEC-2000 column at 30°C. Condition 1: the T4lig2-CVB3-GFP RNA was column purified after T4 ligation. Condition 2: the cleaned-up RNA was heated to 65°C for 3 minutes and immediately placed on ice until injection. Condition 3: T4 RNA ligase buffer (NEB) was added to the cleaned-up RNA sample, containing 50 mM Tris-HCl, 10 mM MgCl₂, and 1 mM DTT, pH 7.5. Condition 4: the column-purified RNA was mixed with T4 RNA ligase buffer before snap cooling. The main and spliced-out intron fractions, marked in blue and red, respectively, were analyzed by the 2% non-EX gel electrophoresis. **(B)** Conditions as in (A) but using column-purified PIE-CVB3-GFP RNA samples and analyzed by 3.5% urea PAGE. Two adjacent intron HPLC fractions were combined and loaded into a single lane for PAGE analysis (lanes 8-10), while the main HPLC fraction was analyzed as single fractions per lane (lanes 1-7).

### HPLC-SEC purifies circRNAs without the need for prior RNA cleanup

We explored whether the RNA cleanup step prior to HPLC-SEC purification could be omitted. Based on the sample pre-treatment tests above, we reasoned that the magnesium ions present from IVT and circularization reaction hindered circRNA separation in SEC. Consistent with our expectations, the circRNAs from both the PIE and STS constructs exhibited poor separation at a column temperature of 30°C (Figure 6A, S6A). The chromatographic profiles of unpurified PIE and STS IVT products nearly mirrored those of spin column-purified RNA samples treated with magnesium and snap cooling (Figure 5B, S5, Condition 4), where introns and circRNAs co-fractionated with precursors. Raising the column temperature to 45°C enhanced separation, with circRNAs and spliced-out introns appearing in later peaks. At 60°C, the separation resolution improved further, closely resembling spin column-purified RNA samples, in which only minimal intron retention was detected in the circRNA fractions (Figure 6A, S6A). To remove the remaining trace of retained introns in PIE-CVB3-GFP circRNA fractions, we combined the strategies of snap cooling and the addition of EDTA and/or formamide (Figure 6B). With 10mM EDTA and 50% formamide present in the samples before injection, the intron content in the circRNA fractions was significantly reduced, and snap cooling further eliminated intron retention to an undetectable level, even in the overexposed PAGE gels (Figure 6D). Additionally, we observed an increased signal of the nicked RNA peak in PIE samples treated with EDTA, formamide, and snap cooling (Figure 6B-C). The high-molecular-weight (HMW) aggregates were dissolved in the presence of EDTA/formamide and heating conditions, leading to an increased nicked circRNA signal. Meanwhile, removing introns also contributed to the decreased UV signals in circRNA fractions. However, this synergistic benefit of formamide/EDTA and snap cooling for circRNA separation was not observed in the STS construct, where all intron products were already removed from both precursor and circRNA fractions at a column temperature of 60°C (Figure S6).

**Figure 6.**
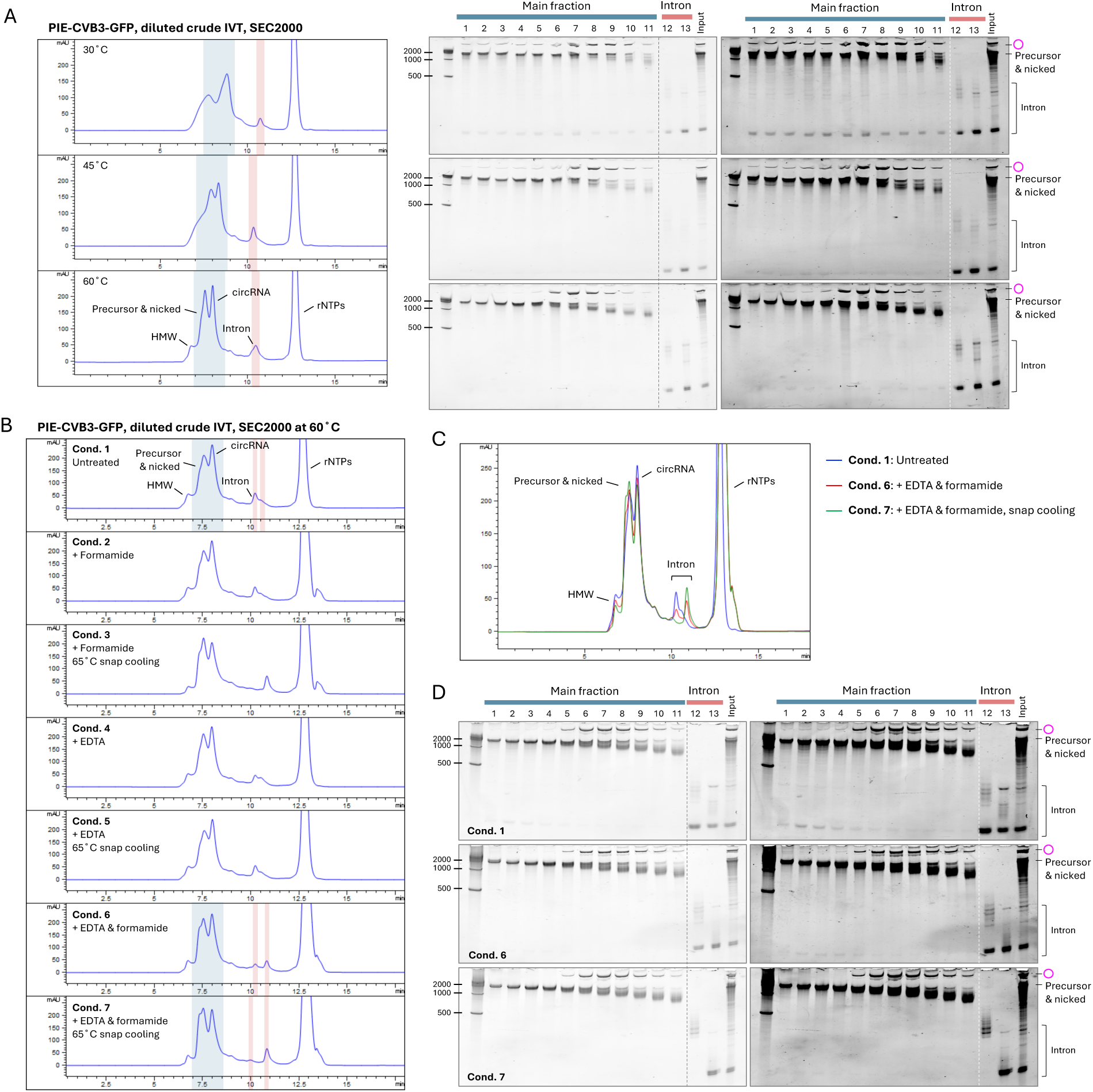
HPLC-SEC purification of PIE-CVB3-GFP circRNAs without prior RNA cleanup. **(A)** HPLC-SEC purification of unpurified PIE-CVB3-GFP IVT products at different column temperatures. Unpurified PIE-CVB3-GFP IVT products (20 µL) were treated with DNase I, circularized by incubation with 2.5 µL 10x T4 ligase buffer (NEB) and 1 µL GTP (100 mM) in 50 µL at 55°C for 30 minutes, and diluted to 100 µL with MEGAclear elution buffer (0.1 mM EDTA, pH 8.0). A 2 µL aliquot was adjusted to 10 µL with the MEGAclear elution buffer and subjected to HPLC-SEC purification using the SEC-2000 column at 30°C, 45°C, or 60°C. The main and spliced-out intron fractions, shown in blue and red, respectively, were analyzed by 3.5% urea PAGE. Gels were visualized using low- and high-contrast settings (left and right images, respectively). **(B)** Effect of sample pre-treatment on HPLC-SEC purification. Following the same sample preparation steps as in A, samples were pretreated using the indicated conditions before injecting them into the SEC-2000, column heated to 60°C. The following pre-treatment conditions were conducted: Condition 1: no additives. Condition 2: 50% formamide with heating and snap cooling (65°C for 3 minutes, then on ice). Condition 3: same as Condition 2 but without heating. Conditions 4: same as Condition 2 but adjusted to 10 mM EDTA flowed by heating and snap cooling (as above). Condition 5: same as Condition 4 but without heating. Conditions 6: adjusted to 10 mM EDTA and 50% formamide, followed by heating and snap cooling. Condition 7: Same as Condition 6 but without heating. **(C)** Overlaid chromatogram of Conditions 1, 6, and 7 from (B). **(D)** Samples of Conditions 1, 6, and 7 were analyzed by urea PAGE, using low and high contrast settings (left and right panel, respectively).

One possible explanation is that PIE samples were supplemented with additional magnesium, followed by a heating process at 55°C for 15 to 30 minutes to improve the yield of the circularization, which may require harsher denaturing conditions to separate circRNAs from intron products. In contrast, no extra magnesium or heating step was introduced for STS constructs after IVT. Additionally, the sequence difference between the group I intron species and the construct design could also affect the intermolecular binding affinity between the excised intron and the circRNA (Figure 1B).

## DISCUSSION

In this study, we investigated the fundamental principle for separating in vitro synthesized circRNAs from linear side-products generated from three synthesis methods. We demonstrated that the denaturation condition is crucial for effective circRNA separation in both gel electrophoresis and HPLC size exclusion chromatography.

Under native conditions, RNA molecules adopt complex structures. During gel electrophoresis, the negatively charged RNA molecules are propelled through a gel matrix by the electric field. RNA molecules of different shapes and hydrodynamic sizes experience varying degrees of drag and retention as they thread through the gel pores, leading to differences in their electrophoretic mobility. Non-denatured linear and circular RNAs of identical sequences are similar in hydrodynamic size, making them difficult to separate in native gels. However, circRNAs probably can form overall more compact structures due to their circular topology, allowing them to migrate slightly faster than their linear counterparts during electrophoresis. In size exclusion chromatography, the migration of RNA molecules is driven by pressure-induced liquid flow. Here, the difference in migration speed is primarily determined by the average hydrodynamic size of RNA molecules. Large molecules bypass the porous stationary phase, traveling fast, while small molecules enter the pores more frequently, leading to delayed elution (29). Like native gel electrophoresis, in SEC, linear RNAs presumably form less compact structures exhibiting larger average hydrodynamic sizes compared to circRNAs. Consequently, linear RNAs are eluted earlier than their circular counterparts. However, such difference in average hydrodynamic sizes induced by circular topology appears to be inadequate for effective circRNA separation under native conditions (Figure 7, left panel).

**Figure 7.**
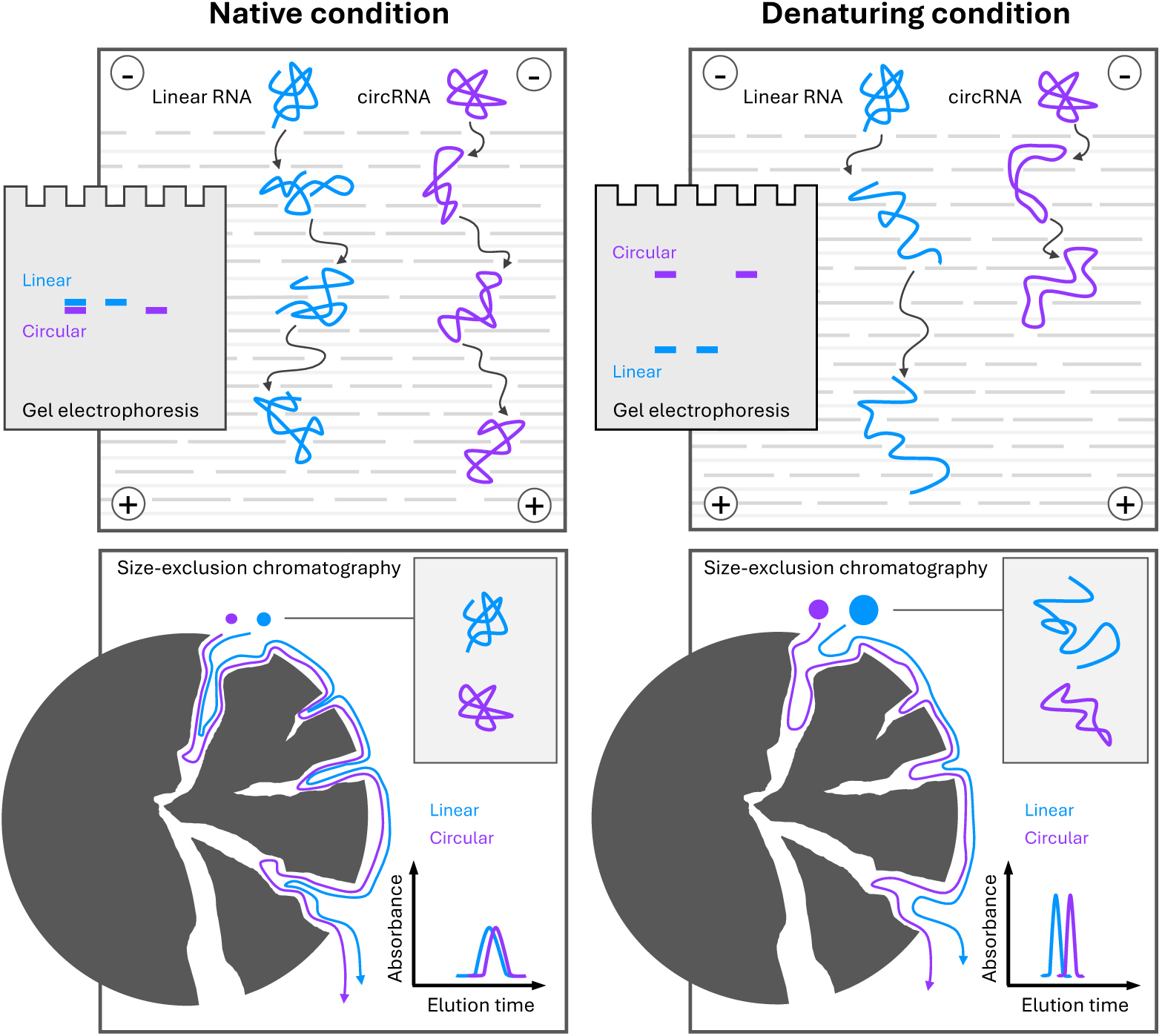
Mechanism of circRNA separation.

Under denaturing conditions, such as high temperature or chemical denaturants like urea and formamide, the complex structures of RNA molecules are unfolded, induced by the disruption of hydrogen bonds. During denaturing gel electrophoresis, linear RNAs are likely forced into an extended single-stranded configuration, allowing them to navigate the gel matrix by “snaking” through the pores. In contrast, circRNAs still stay in a looped structure, resulting in increased friction as they may traverse the gel pores in a manner resembling unpaired double-stranded RNA molecules. This leads to markedly slower migration and distinct banding patterns compared to their linear counterparts. Considering the agarose E-Gel system, for which the gel components and the electrophoresis parameters are proprietary and undisclosed, we assume that the excessive heat buildup during electrophoresis is denaturing the RNA, consequently facilitating circRNA separation. In size exclusion chromatography, under denaturing conditions, both linear and circular RNAs become “floppier” and more expanded than their native conformations, leading to larger average hydrodynamic sizes and generally shorter elution times. Importantly, due to the structural constraints of a circular topology, even when fully denatured, circRNAs can only reach half the maximum length of their stretched linear counterparts. This results in a greater difference in average hydrodynamic sizes between circular and linear RNAs under denaturing conditions, thus improving the efficiency of circRNA separation (Figure 7, right panel). We speculate this principle could extend to CGE for effective circRNA separation by using only formamide as the capillary gel solvent, a method successfully applied in high-resolution linear mRNA analysis (47).

Besides the structural status of RNA molecules, the size of circRNA directly impacts its separation in gel electrophoresis. Our investigation primarily focused on the CVB3-GFP circRNAs of approximately 1500 nt, while circRNAs of varying sizes can exhibit distinct electrophoretic patterns. For instance, small RNA circles (10 to 25 nt) migrate faster than their linear counterparts in 12% to 18% denaturing urea PAGE (43,48,49). The sieving ability of the gel matrix is also strongly influenced by the concentration of agarose or polyacrylamide. Our study revealed that circular and linear CVB3-GFP RNAs of equivalent lengths overlapped in 1% agarose gel electrophoresis, regardless of whether self-cast gels or commercial E-Gels were used. This overlap likely stems from the pore size of the gel matrix being too large, preventing linear RNA molecules from adopting a single-stranded configuration as they migrate through the pores, thereby reducing the separation efficiency for circRNAs. Notably, a recent study showed that 0.8% native agarose gel with extended electrophoresis time successfully separated a super-large RNA circle of 8600 nt from its linear counterpart (10). These findings highlight that the electrophoretic separation of circRNAs depends on the interplay between RNA size and the sieving effect of the gel matrix.

An intriguing observation in our study is that using HPLC-SEC at 45°C with a Tris-EDTA mobile phase improved the separation of T4-ligated and PIE-generated circRNAs at low pH (< 4.5) and neutral pH (7.5), whereas intermediate pH levels (5.5-6.5) showed poorer performance. One possible explanation for better circRNA separation at low pH is that protonation at N3 position in cytosine (pKa 4.2) (50) disrupts the hydrogen bonds in G-C pairs and promotes denaturation of the RNA molecules. Thus, both linear and circular RNAs may become less structured and more homogenous, allowing them to be separated more effectively in HPLC-SEC.

Magnesium ions in RNA samples are an easily overlooked factor because they can co-purify with the RNA. Even a small amount can significantly affect HPLC-SEC performance, leading to intron retention in PIE and STS circRNA fractions. Intron co-elution in circRNA fraction is likely caused by the remaining base pairing of the intronic ribozyme sequences to the guide sequences of the exons (Figure 1A). We found that column heating, combined with the addition of EDTA to chelate magnesium ions and formamide to destabilize hydrogen bonding in the RNA samples, following snap cooling to avoid intermolecular base paring, effectively resolved this issue.

After our extensive efforts to optimize circRNA separation using HPLC-SEC, several limitations remain. (i) Even at optimized conditions, SEC purification cannot fully separate circRNA from smaller side products. Since circRNAs are eluted later than their linear counterparts, circRNA can co-elute with impurities from in vitro transcription or degradation products during HPLC processing. As a result, additional purification steps, including RNase R treatment, are required to increase the purity of circRNAs. This issue is further exacerbated when working with smaller circRNAs, as their fractions may overlap with the spliced-out intron fragments that have similar elution times. (ii) The separation efficiency of circRNAs in SEC is further compromised when injecting larger amounts of RNAs, which hampers upscaling purification (Figure S7). (iii) RNA is susceptible to degradation at high temperatures and shear force exerted by the column matrix, leading to increased nicking of circRNAs. However, according to chromatographic profiles, we did not observe significant loss of circRNA signal when heating the column, likely attributed to the short processing time, typically around 10 minutes from injection to elution in SEC-2000. (iv) Heating the column is discouraged by the manufacturer, as excessive heat can shorten its lifespan and diminish its performance. Clearly, further development of scalable circRNA separation methodologies is still needed.

Already in the 1970s, scientists used denaturing PAGE to successfully separate and analyze in vitro synthesized circRNAs (19,48). This pioneering achievement provided a powerful and straightforward biochemical approach for studying circRNAs, which are commonly associated with self-splicing introns – a prominent life science topic during the 1980s. Research on synthesized circRNAs fell into abeyance around the turn of the 21st century but resurged with the advent of mRNA vaccine. However, the current rapid development of circRNA research has been occasionally accompanied by a diminished focus on fundamental RNA biology knowledge. In our study, we revisit previously established concepts and further explore the principle underlying circRNA separation. We believe our findings will advance the development of circRNA-based biotechnology and therapeutics, paving the way for innovative treatment strategies in the future.

## DATA AVAILABILITY

Data supporting the findings of this study are available within the article and its supplementary material.

## AUTHOR CONTRIBUTIONS

Y.J. performed laboratory experiments and data analysis. Y.J. and J.K. conceived the idea, J.K. supervised the project. Y.J. wrote the draft, J.K. edited and revised the manuscript.

## ACKNOWLEDGEMENT

The authors thank Rasmus Rytter Hartmann for his technical support with HPLC. We also thank Maria Gockert for reading and editing this manuscript and the members of the Kjems lab for all the inspiring discussions. This work was supported by Novo Nordisk Foundation [NNF23OC0081177 and NNF22SA0081890]. Y.J. was funded by the China Scholarship Council [202106010111].

---

**Figure S1.**
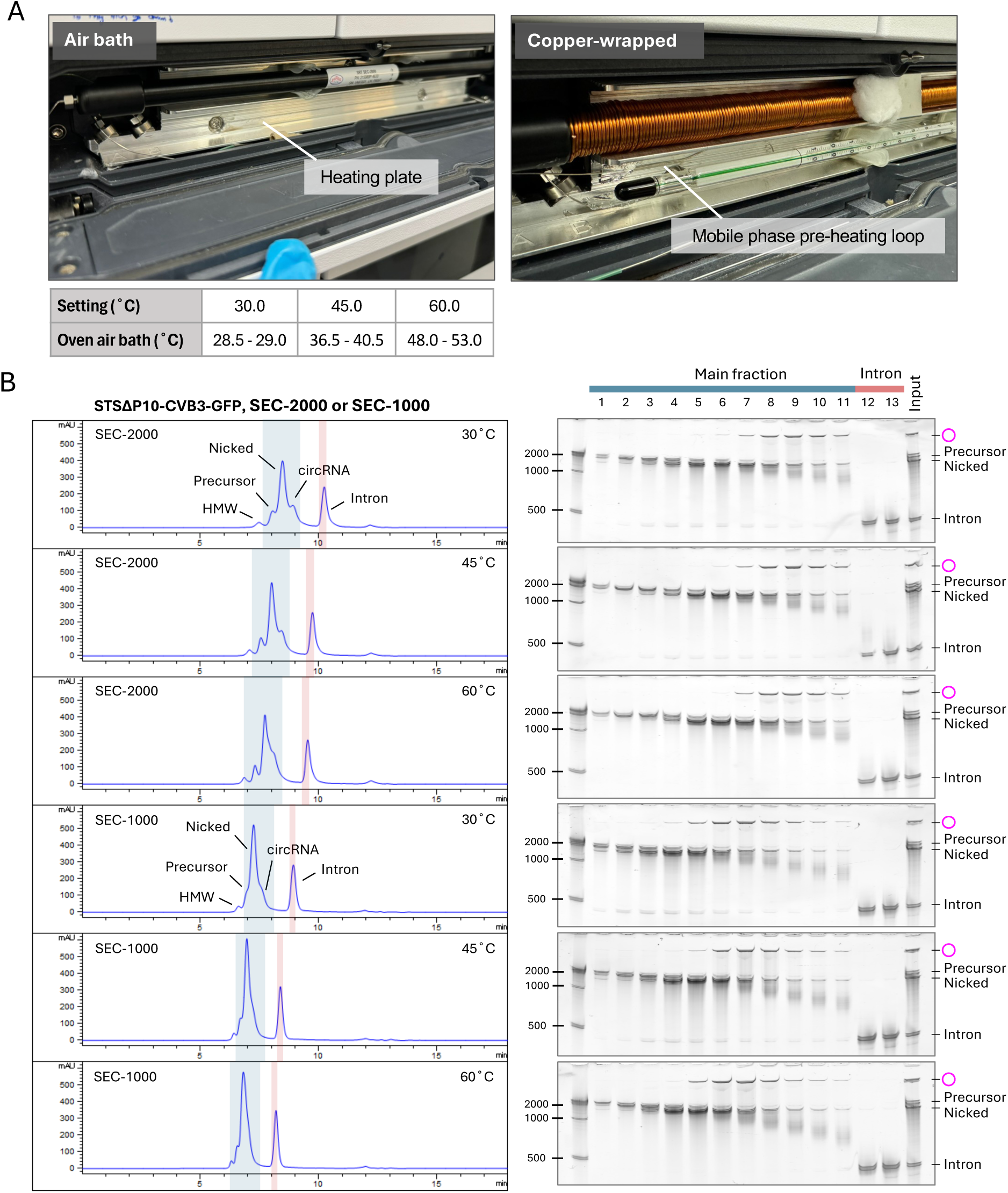
HPLC-SEC column setup and purification of STSΔP10-CVB3-GFP using SEC-1000 or SEC-2000. (related to. **Figure 4) (A)** The HPLC-SEC column setup. The air temperatures in the column oven were measured and compared to the setting temperatures. **(B)** circRNA separation of STSΔP10-CVB3-GFP in SEC-1000/2000 with column heating. RNAs were purified with HPLC-SEC at 30, 45 and 60°C. The eluted fractions were analyzed using 3.5% urea PAGE, with main and intron fractions marked in blue and red.

**Figure S2.**
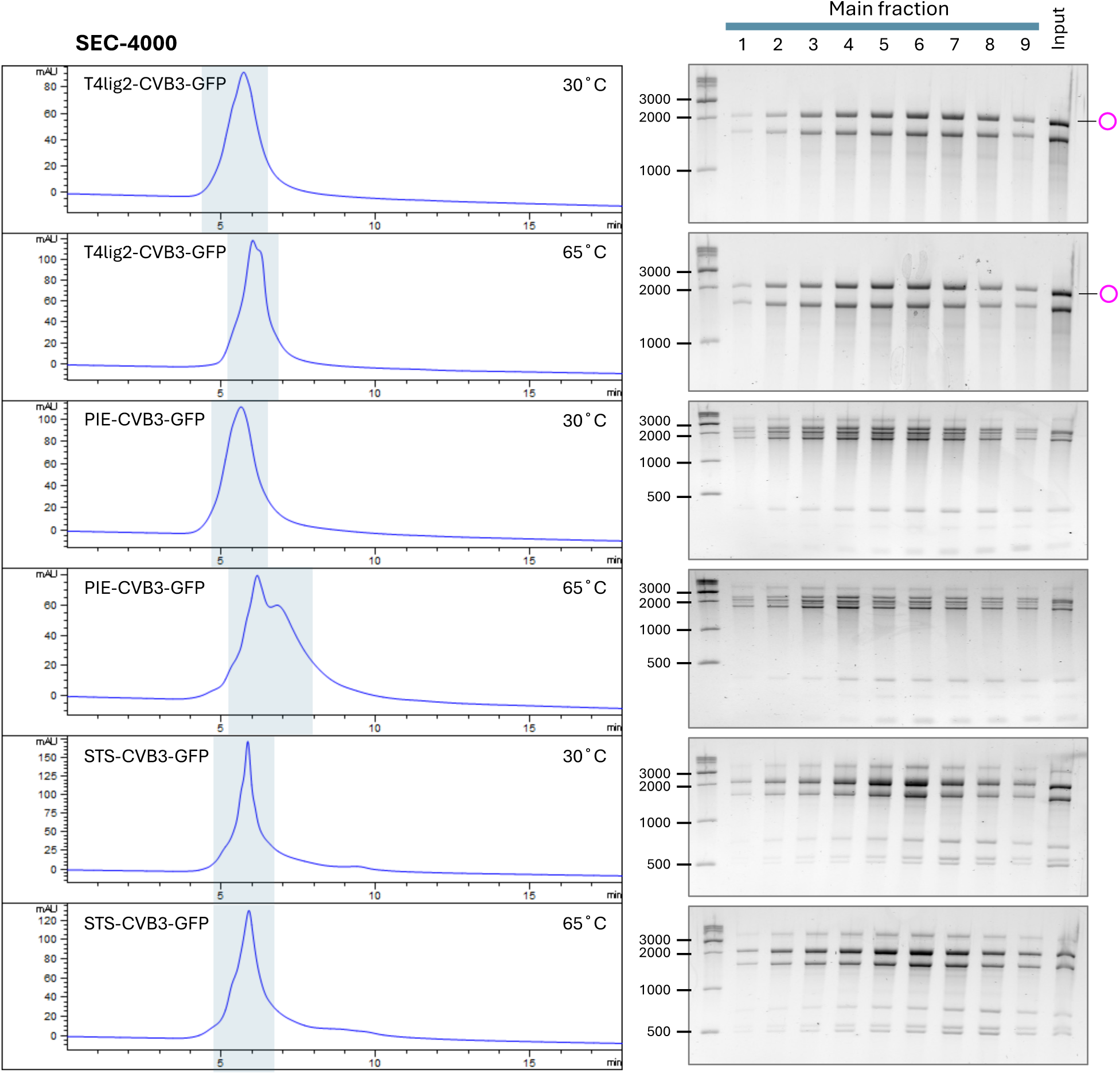
HPLC-SEC purification of circRNAs using SEC-4000. Cleaned-up RNA samples were injected into the SEC-4000 column at 30 or 65°C. The collected fractions (depicted in blue areas) were analyzed with 2% EX gel electrophoresis. The circRNAs are marked with magenta circles to the right.

**Figure S3.**
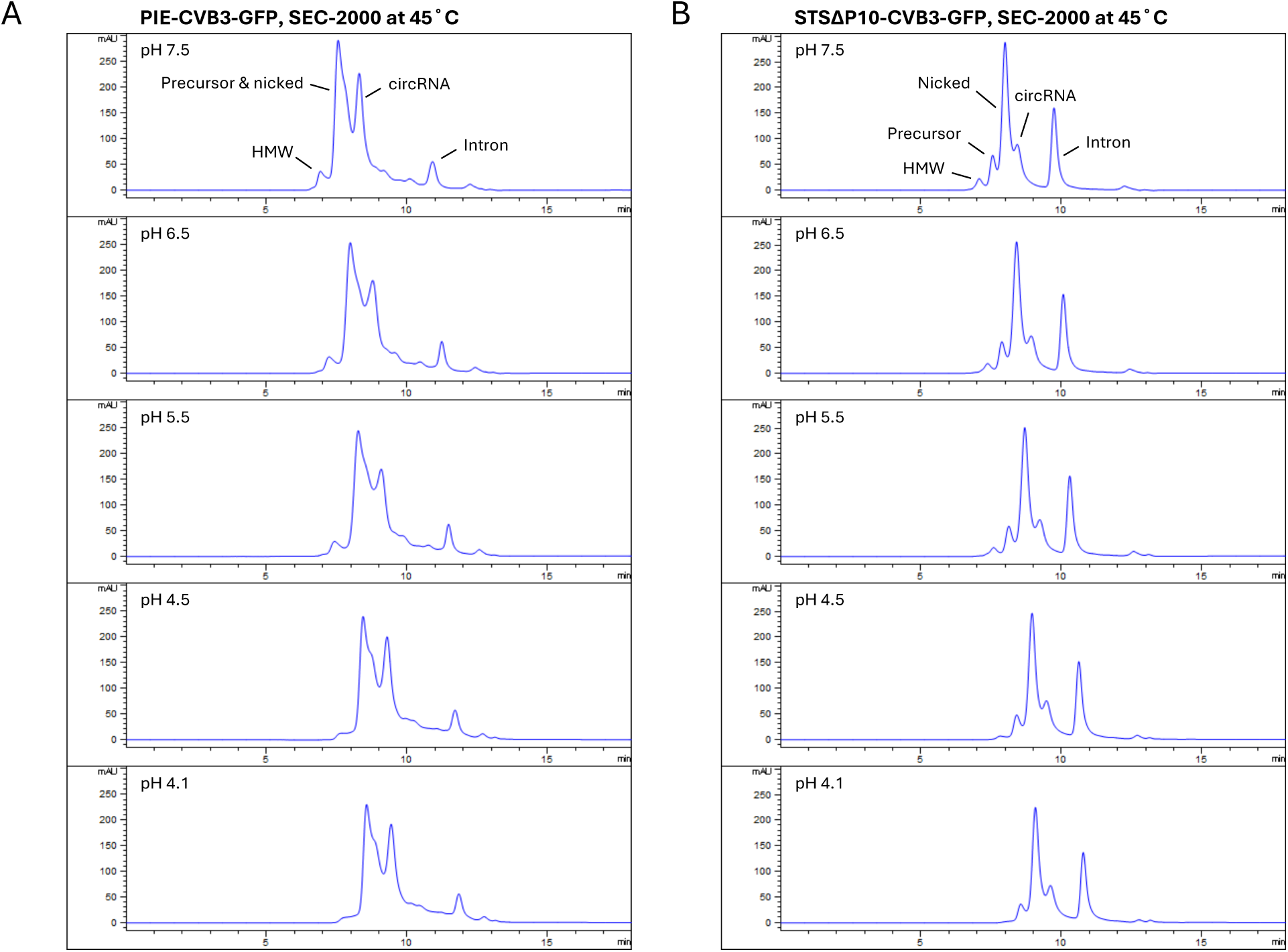
HPLC-SEC purification of ribozymatically synthesized circRNAs using mobile phases of the indicated pH (related to. **Figure 4). (A and B)** PIE- and STSΔP10-CVB3-GFP RNA samples were purified by the SEC-2000 column at 45°C using mobile phases with varying pH.

**Figure S4.**
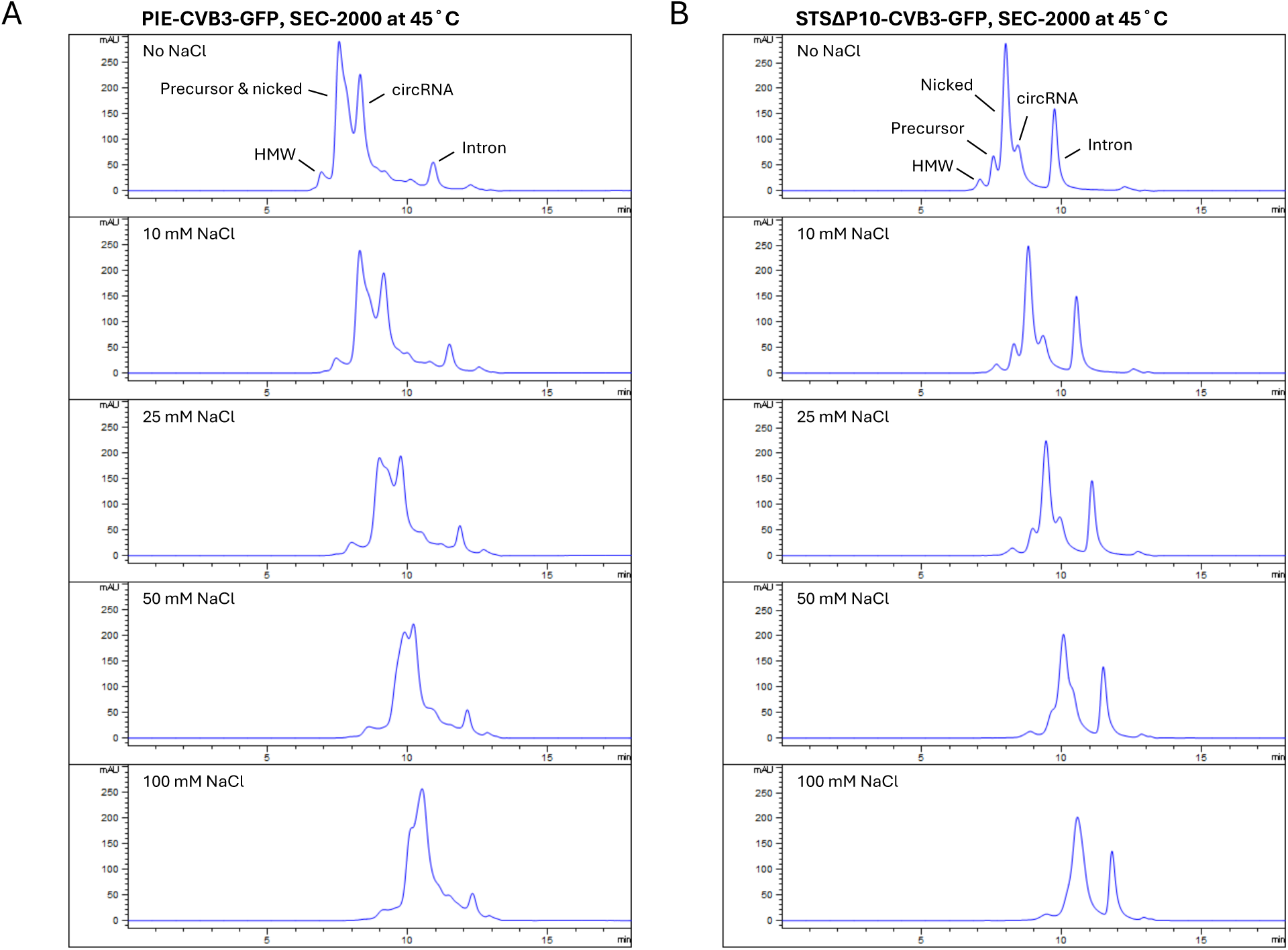
HPLC-SEC purification of ribozymatically synthesized circRNAs with mobile phases containing the indicated sodium chloride concentrations (related to. **Figure 4). (A and B)** PIE- and STSΔP10-CVB3-GFP RNA samples were purified by the SEC-2000 column at 45°C with mobile phases of different salt concentrations.

**Figure S5.**
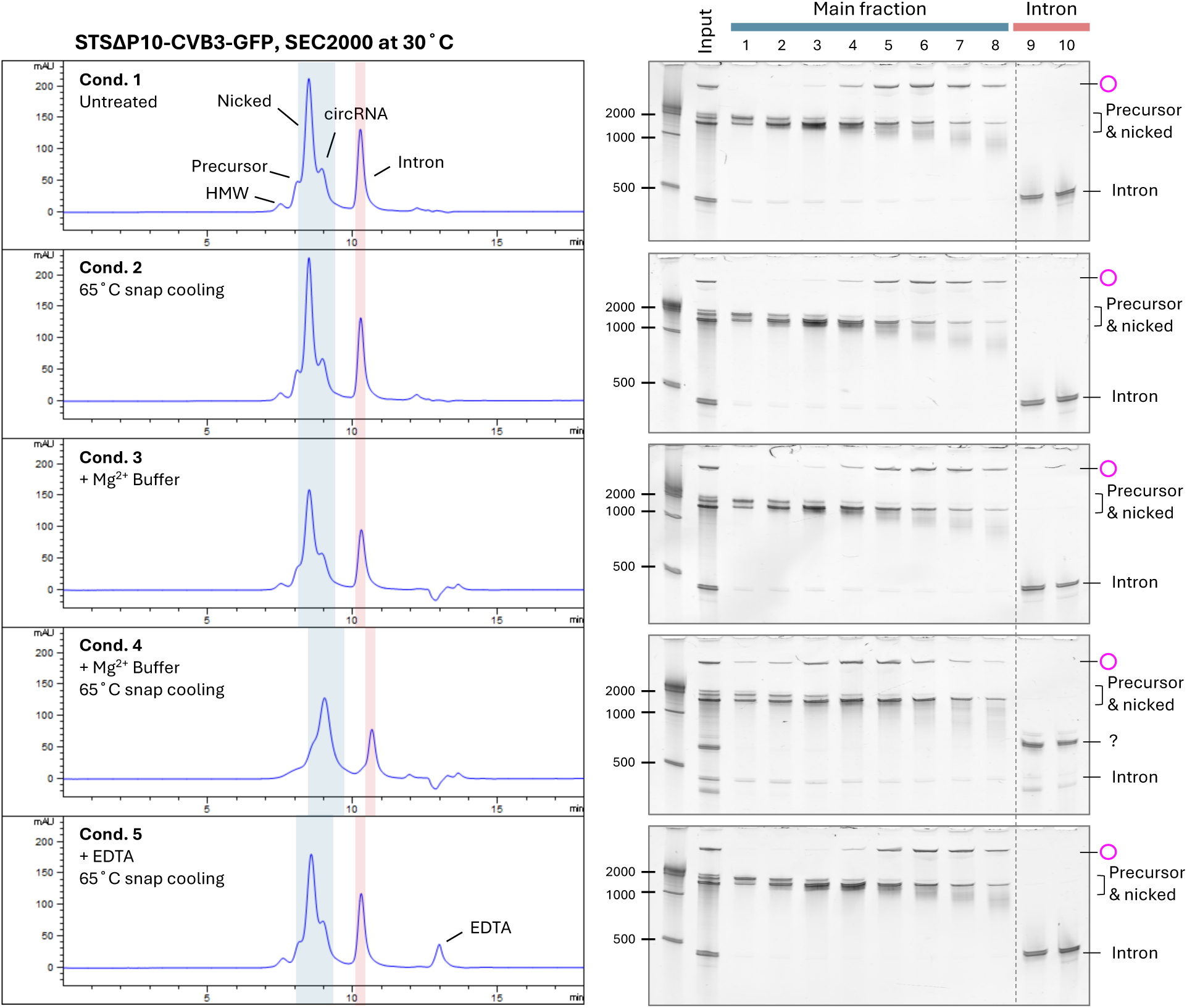
HPLC-SEC purification of STSΔP10-CVB3-GFP RNA with the indicated sample pre-treatment conditions (related to. **Figure 5).** Column-purified STSΔP10-CVB3-GFP RNA samples were processed as in Figure 5, except for the last condition. Condition 5: EDTA (10 mM) was added to the cleaned-up RNA, followed by snap cooling. HPLC-SEC fractions were analyzed by 3.5% urea PAGE. The analyzed main and intron fractions are shown in blue and red.

**Figure S6.**
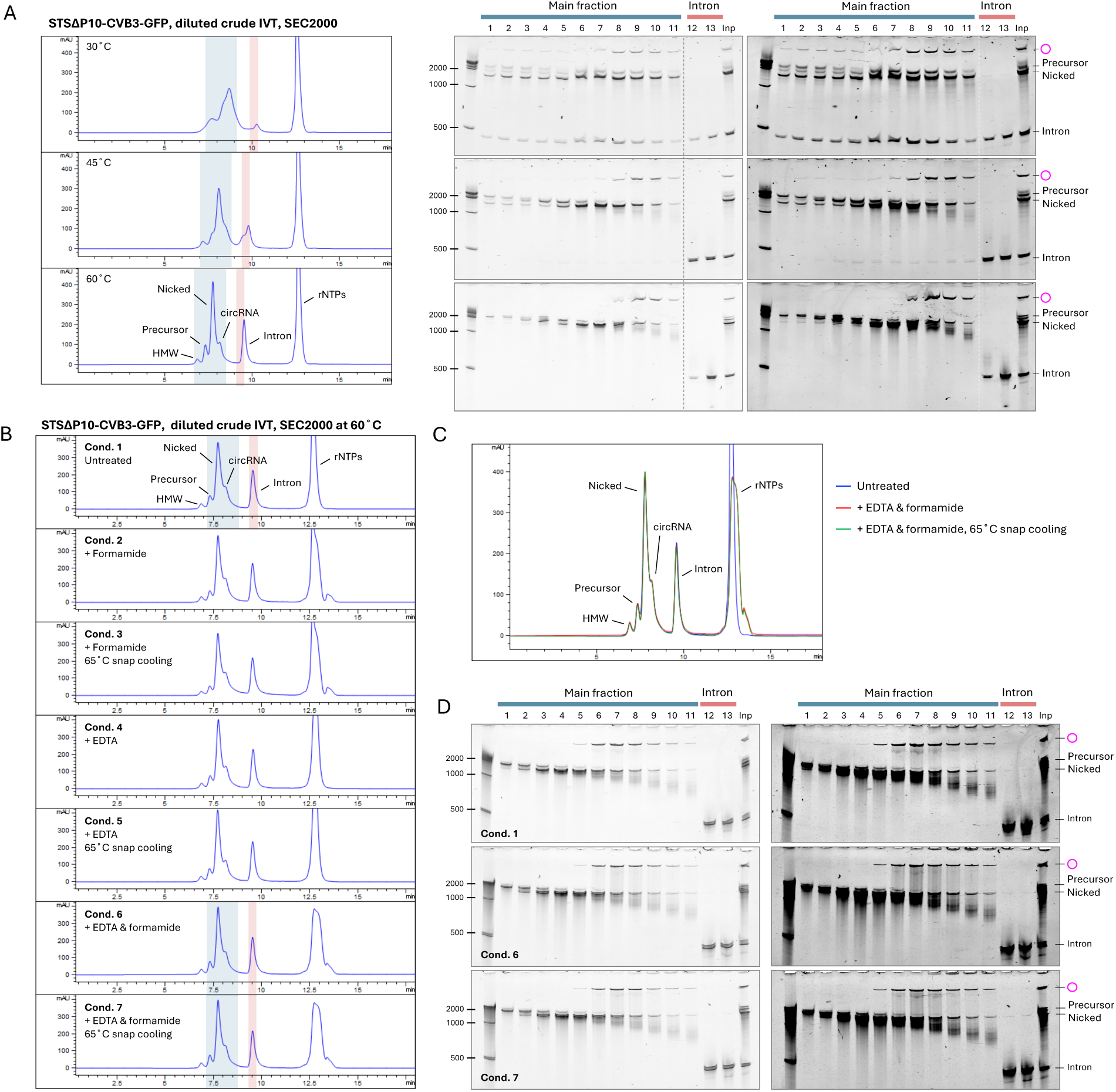
HPLC-SEC purifies STSΔP10-CVB3-GFP circRNAs without prior RNA cleanup (related to. **Figure 6). (A)** HPLC-SEC purification of diluted crude STSΔP10-CVB3-GFP IVT products using the SEC-2000 column at 30°C, 45°C, and 60°C. After IVT and DNase treatment, the crude product (50 µL) was diluted with MEGAclear elution buffer (0.1 mM EDTA, pH 8.0) to 100 µL. A 2 µL aliquot was further diluted to 10 µL using the same buffer, followed by HPLC-SEC purification. The main and intron fractions marked in blue and red, respectively, were analyzed by 3.5% urea PAGE. Gels were visualized using low and high contrast settings. **(B)** Pre-treatment of STSΔP10-CVB3-GFP IVT prior to HPLC-SEC, following the same procedure as in Figure 6B. The main and intron fractions, shown in blue and red, were analyzed by 3.5% urea PAGE. **(C)** Chromatogram overlay of the three pre-treatment sets from (B). **(D)** Samples of conditions 1, 6, and 7 from (B) were analyzed by 3.5% urea PAGE, visualizing gels using low and high contrast settings.

**Figure S7.**
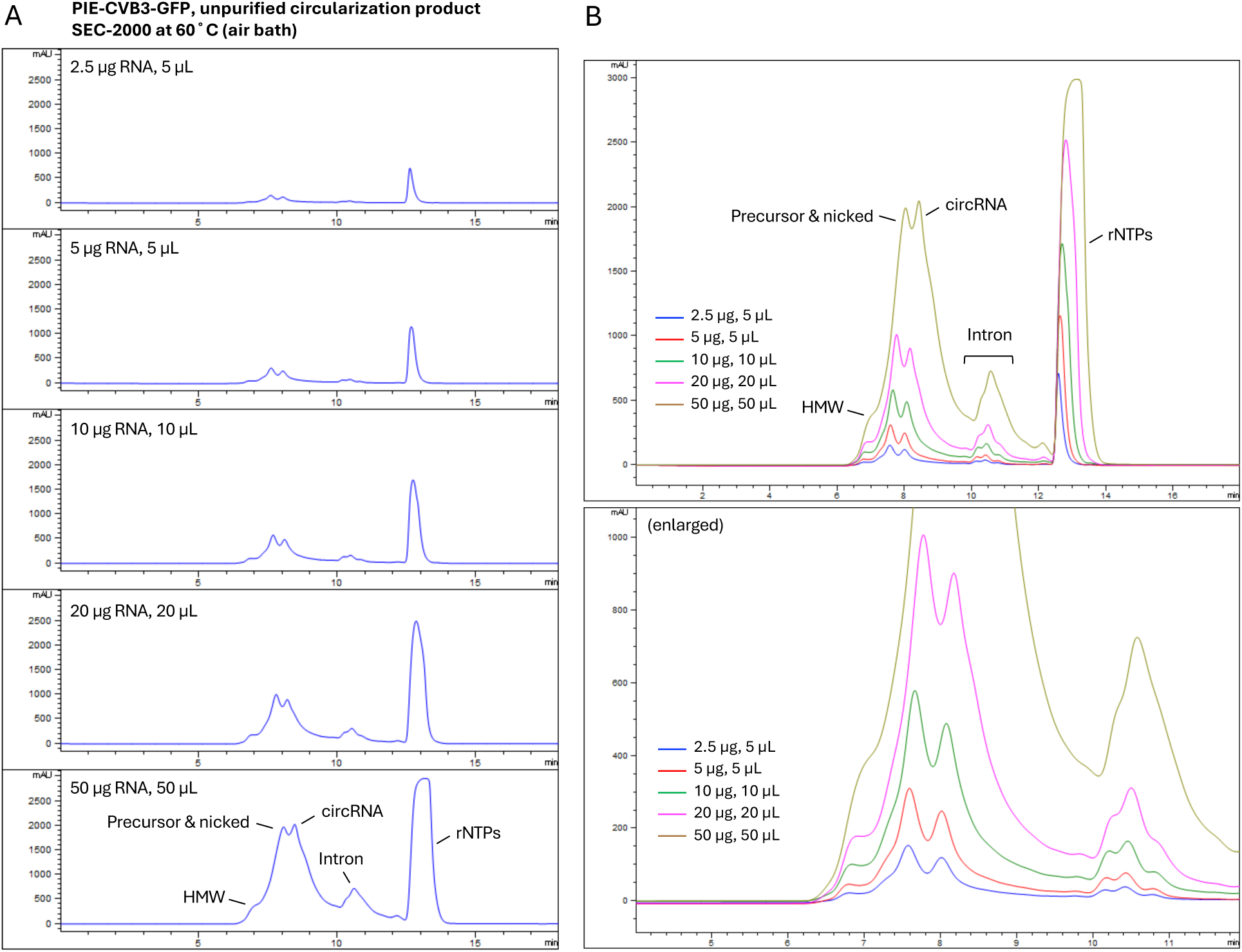
RNA loading capacity of SEC-2000 column. **(A)** HPLC-SEC purification of the PIE-CVB3-GFP unpurified circularization product using SEC-2000 column at 60°C (air bath). After IVT and DNase treatment, the RNA product was purified by a spin column. Then, T4 RNA ligase buffer (NEB) and GTP (2 mM) were added, followed by a 15-minute incubation at 55°C. The unpurified reaction was directly injected into HPLC-SEC with varying RNA amounts and injection volumes. **(B)** Chromatographic overlay shown in (A).

**Figure S8.**
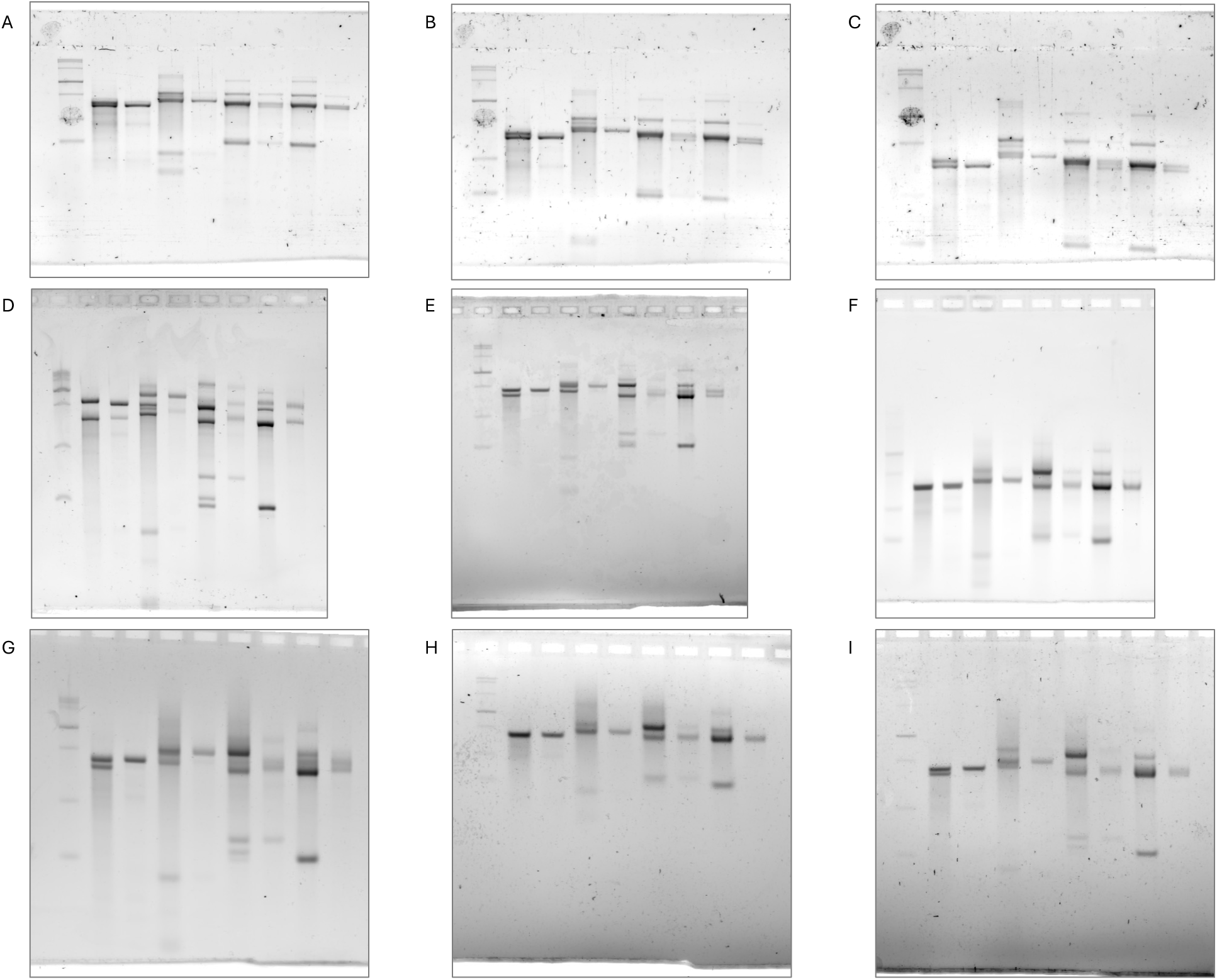
Unprocessed gel images from. **Figure 2. (A to C)** Unprocessed gel images from Figure 2C. **(D to F)** Unprocessed images from Figure 2D. **(G to I)** Unprocessed images from Figure 2E.

**Figure S9.**
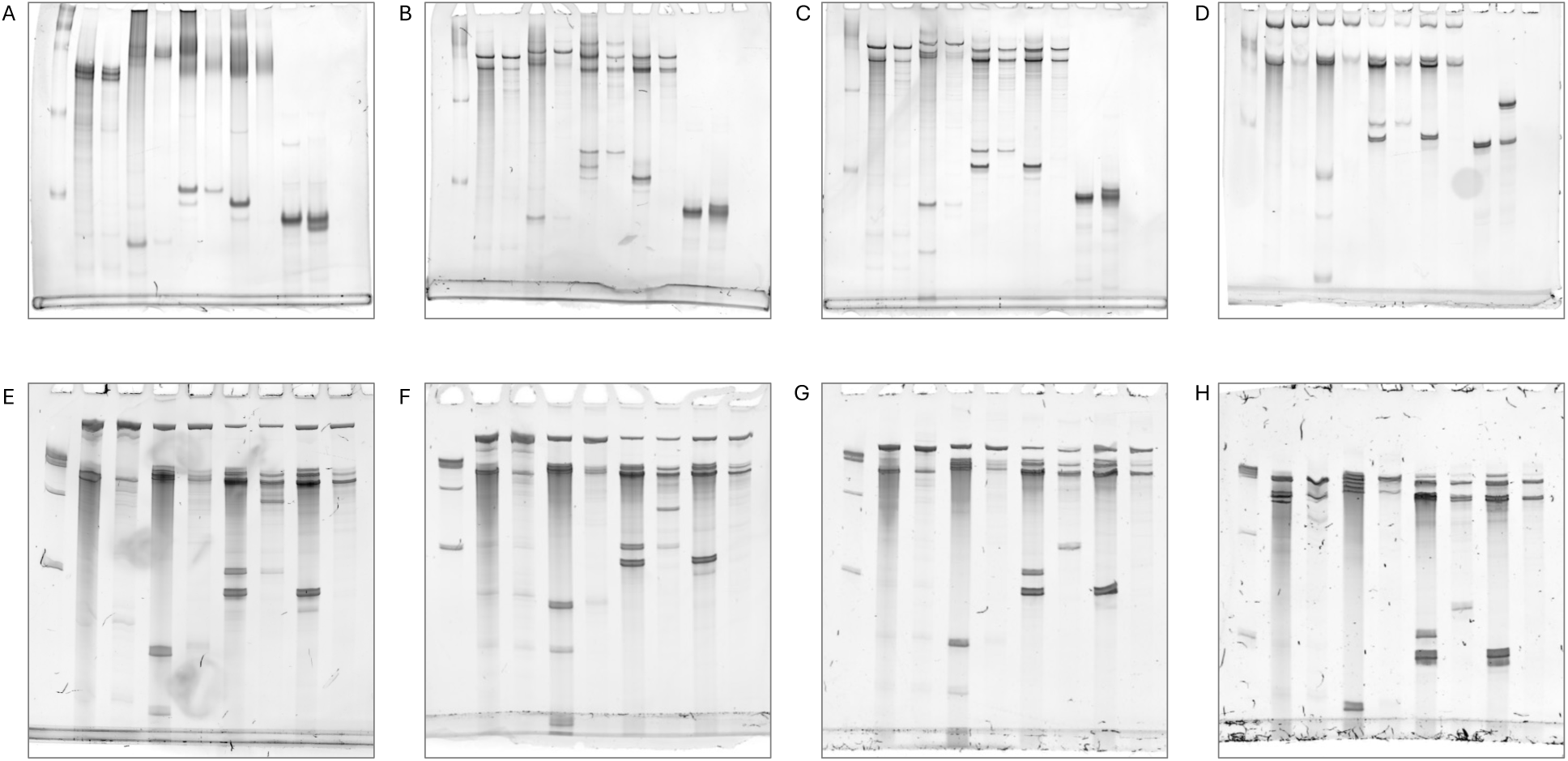
Unprocessed gel images from. **Figure 3. (A to D)** Unprocessed gel images from Figure 3A. **(E to H)** Unprocessed gel images from Figure 3B.

**Table S1.**
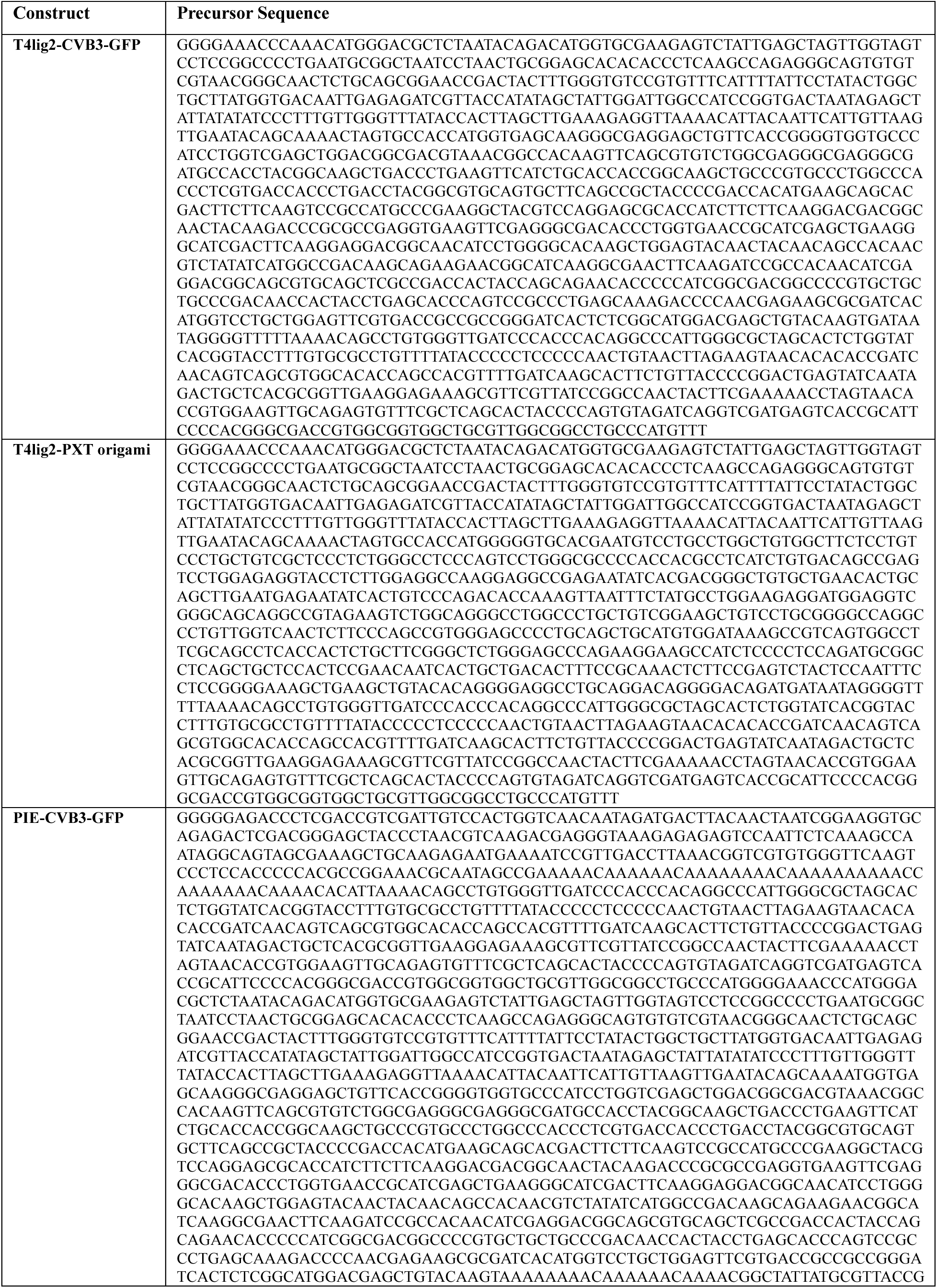
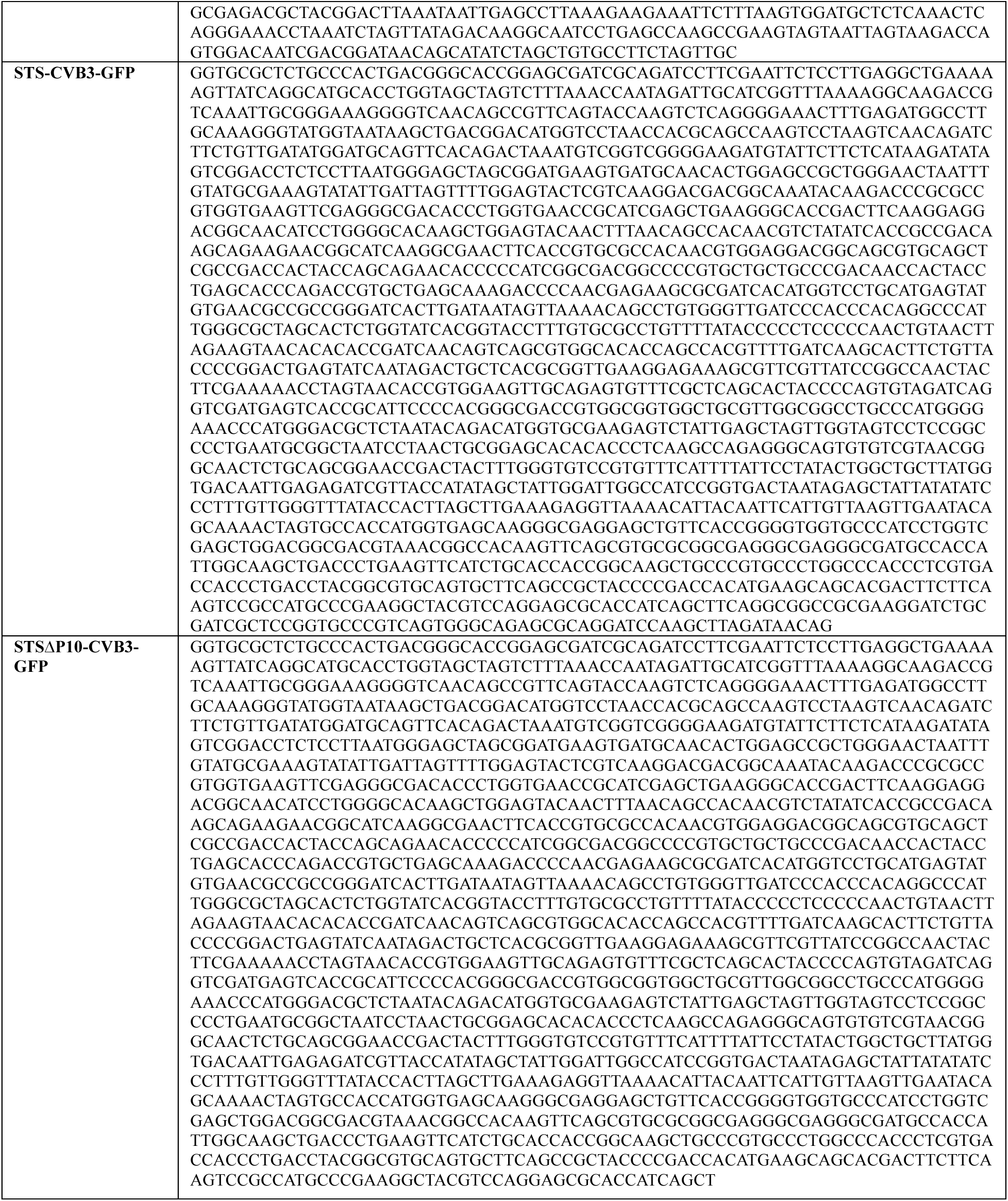
The transcribed precursor sequence.

